# Deep Profiling of EV Long RNAs Reveals Biofluid-Specific Transcriptomes and Splicing Landscapes

**DOI:** 10.1101/2025.08.05.668716

**Authors:** Sudipto K. Chakrabortty, Shuran Xing, Sinead Nguyen, Allan George, Dulaney L. Miller, Kailey Babcock, Kyle Manning, T. Jeffrey Cole, Emily Mitsock, Christian J. Ray, Sivakumar Gowrisankar, Johan Skog

## Abstract

RNA profiling of extracellular vesicles (EVs) from human biofluids has historically been limited to small RNA species, with long RNAs—such as mRNA exons and long non-coding RNAs—remaining largely underexplored. Moreover, the dominance of hematopoietic-derived EVs in complex fluids like plasma has posed significant challenges for detecting low-abundance, tissue-specific transcripts. Here, we establish foundational transcriptomic maps of long RNAs in EVs from plasma, urine, and cerebrospinal fluid (CSF) using ultra-deep whole transcriptome sequencing (WTS), revealing both fluid-specific and shared expression and splicing signatures. We then introduce a targeted RNA capture method that enriches for all protein-coding and long non-coding transcripts, dramatically enhancing sensitivity for gene and splice variant detection. Applying this approach to brain-specific transcripts, we achieve >85-fold enrichment of target gene expression and, on average, 3.1-fold increase in detected splice junctions per gene compared to untargeted WTS. As a proof of concept, we apply this brain-targeted RNA panel to EVs from plasma in a Parkinson’s disease cohort of 40 plasma samples and compare its performance to exome sequencing as well as untargeted WTS. This work advances EV transcriptomics into the long RNA domain and establishes a framework for high-sensitivity, noninvasive biomarker profiling across tissues and diseases.

## Introduction

Liquid biopsy approaches, including analysis of circulating cell-free DNA (cfDNA), circulating tumor cells (CTCs), and extracellular vesicles (EVs), have transformed noninvasive biomarker discovery, with demonstrated clinical utility in cancer and other diseases[1–5]. EVs offer unique advantages as molecular messengers, carrying RNA, DNA, proteins, and lipids from their cell of origin [1, 6–8]. EVs are highly stable and reflective of active cellular processes, making them attractive vehicles for biomarker development across a broad range of pathologies, [9–12], including cancer [13–22], neurodegenerative disorders [23–31], cardiovascular diseases [32–39], and immune dysfunction [40–46].

Most transcriptomic studies of EV RNA have focused on small non-coding RNAs (e.g., microRNAs, tRNAs, and Y RNAs) [47–52]. In contrast, the landscape of long RNAs—such as mRNAs and long non-coding RNAs—remains less characterized [53–56]. While whole transcriptome sequencing (WTS) has revealed the broad diversity of long RNAs in EVs, its sensitivity is limited by the dispersion of reads across genomic regions of low relevance. Furthermore, the complexity of the EV transcriptome, particularly at the level of isoform diversity and alternative splicing, remains largely unexplored.

Targeted RNA sequencing using hybrid capture can overcome these limitations by enriching for protein-coding and long non-coding transcripts, enabling deeper profiling of gene expression, RNA splicing, and fusions. This is particularly valuable when analyzing EVs from complex biofluids such as plasma, where hematopoietic-derived vesicles dominate and dilute signals from tissue-specific transcripts [9, 57–59]. Although surface-marker-based enrichment strategies for tissue-specific EVs are under investigation, their practical utility remains limited [60–63].

Here, we present a comprehensive analysis of EV long RNAs across three human biofluids— plasma, urine, and cerebrospinal fluid (CSF)—using both WTS and a targeted hybrid-capture approach (Whole Exome RNA Sequencing, WES). We map the fluid-specific and shared transcriptomic profiles with unprecedented depth, revealing fluid-enriched transcripts and extensive isoform diversity. Leveraging brain-enriched RNA capture panels, we demonstrate highly sensitive detection of rare, neuronal-derived transcripts and splice variants from plasma EVs. As a proof of concept, we apply these approaches to a Parkinson’s disease (PD) cohort and compare the performance of WTS, WES, and brain-targeted sequencing for biomarker discovery. Finally, we evaluate the added value of integrating gene expression and splicing signatures, establishing a framework for multiomic EV-RNA liquid biopsy diagnostics.

## Results

### EV RNA Profiling of Biofluids by Whole Transcriptome Sequencing

We began by performing deep sequencing of EV RNA from plasma, CSF, and urine, obtained from healthy donor pools by WTS, which showed notable distinctions in alignment, read distribution and biotypes (Figure 1). Plasma samples consistently showed the highest proportions of read alignment, with over 80% of the reads aligned to the human genome (Figure 2a). In contrast, CSF samples consistently showed the lowest read alignment, with over 60-70% of reads unable to be mapped to the human genome. Urine samples, on the other hand, demonstrated large sample-to-sample variability, with read alignment varying from 40 to 80%. Next, we investigated the genomic origin of reads in biofluid-derived EVs. We observed that the proportion of exonic reads in plasma ranges between 40% - 45% of the total reads (Figure 2b). In contrast, the majority of reads in CSF samples remained unmapped with less than 15% reads mapping to exons, suggesting low quality and quantity of input RNA. Urine samples continued to demonstrate high sample-to-sample variation. While two of the urine samples demonstrated reasonably high exonic mapping (45 - 55%), the other two demonstrated intermediate levels of exonic reads. A further investigation on the biotype distribution of transcriptome-mapping reads revealed urine samples demonstrated the highest proportions of reads mapping to protein-coding genes and the most efficient depletion of ribosomal RNA (rRNA) compared to other samples (Figure 2c). Despite superior exome mapping among biofluids, plasma samples showed intermediate levels of protein-coding mapping reads ranging between 50 - 85%, demonstrating sample to sample variability. Interestingly, plasma also demonstrated the highest proportions of small non-coding RNAs (ncRNA) among the biofluids. Despite the poor mappability observed in CSF samples described above, they demonstrated intermediate levels of protein coding reads (40 - 80%). Also of interest, CSF EVs demonstrated the largest proportion of reads mapping to long non-coding RNAs (lncRNAs) among biofluids. The external RNA spike-in controls (ERCC) were generally low in abundance and consistent in all samples except for CSF, indicating that CSF has the lowest RNA yield recovered among the biofluids.

**Figure 1.**
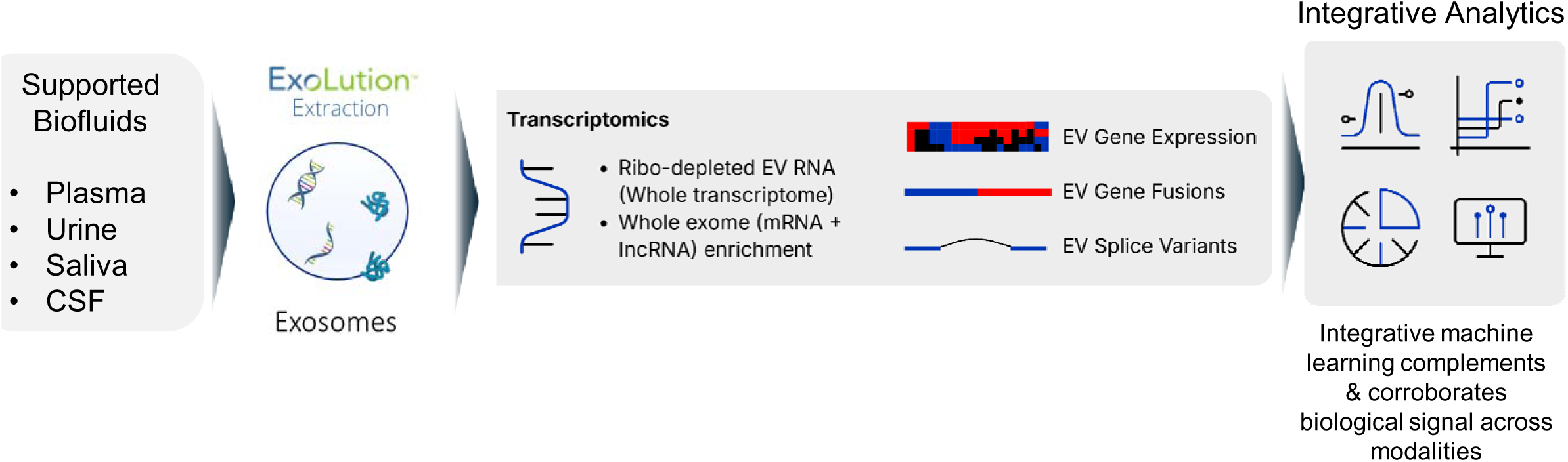
Overview of long RNA profiling from biofluid-derived exosomes.

**Figure 2.**
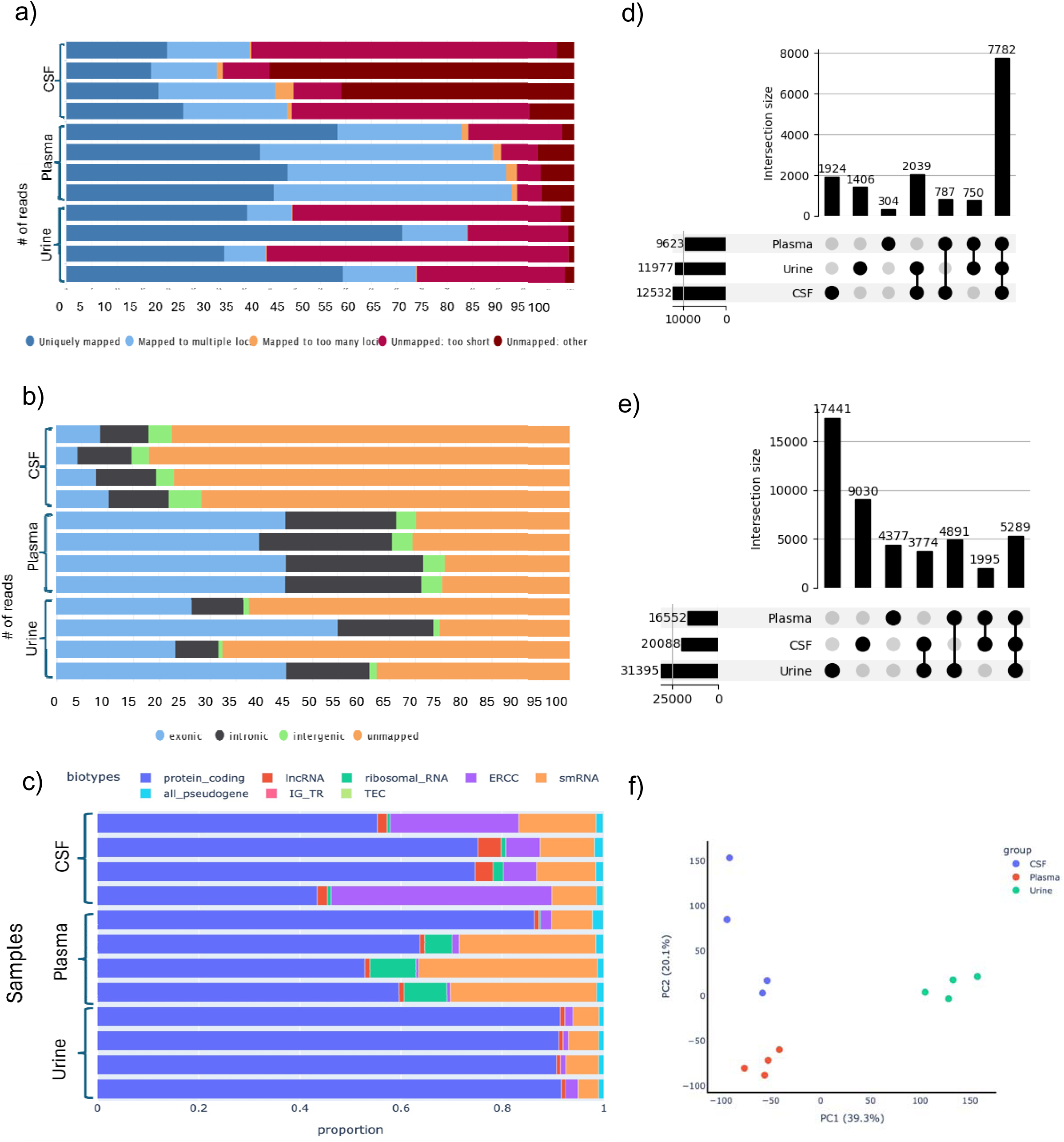
Whole Transcriptome Sequencing across Plasma, Urine, & CSF Samples. a) STAR alignment statistics for plasma, urine, & CSF EV samples. The x-axis is the percentage of reads. The percentage of uniquely mapped reads varied between biofluids. b) Genomic origin of reads for plasma, urine, & CSF EV samples. The x-axis is the percentage of reads. The percentage of reads mapping to exonic regions is less than 50% in most samples. c) Biotype distributions for plasma, urine, & CSF EV samples. The x-axis is the percentage of reads. The majority of the exonic reads map to protein coding regions in all samples. d) Gene detection rate in different biofluid derived EVs with a detection threshold of >1 average TPM across samples. e) Transcript detection rate in different biofluid derived EVs with a detection threshold of >1 average TPM across samples f) Principal component analysis (PCA) shows distinct clustering of each biofluid derived EVs.

Surprisingly, detailed analysis of the protein coding reads revealed that the highest number of protein coding genes across biofluids was detected in CSF (12,532 genes) with more than 1 transcript per million (TPM), followed by urine which detected 11,977 protein coding genes (Figure 2d). Plasma samples resulted in detection of 9,623 protein coding genes, respectively. This is surprising because CSF samples showed the lowest proportion of exonic reads among biofluids (Figure 2b). A comparison of the protein coding gene composition across biofluids revealed that 14,992 protein coding genes were detected from all biofluids in total, of which 7,782 genes were shared among all of them, accounting for 52% of total protein coding genes detected in these biofluids (Figure 2d). Interestingly, CSF EVs demonstrated the highest number of protein coding genes uniquely detected in any biofluid (1,924 genes), followed by urine with 1,406 uniquely detected genes. Plasma samples showed only 304 uniquely detected protein coding genes. Next, we investigated the detection of protein coding transcripts across biofluids. While CSF resulted in the highest number of protein coding genes across biofluids, urine samples resulted in the highest number of protein coding transcripts (31,395 transcripts). This was followed by CSF with 20,088 protein coding transcripts and plasma with 16,552 protein coding transcripts (Figure 2e). The total number of protein coding transcripts detected in all biofluids combined was 46,797, accounting for roughly 28% of the human transcriptome. Comparative analysis across biofluids further revealed that only 5,289 protein coding transcripts (11% of the total no. of transcripts, i.e., 46,797) were detected in all three biofluids. This is in sharp contrast with genes where 52% of genes were co-detected in all biofluids, suggesting biofluid specificity of EV cargo is more pronounced at the transcript level than the gene level. Over 17,400 transcripts were uniquely detected in urine, accounting for ∼55% of all transcripts detected in urine EVs. 9,030 protein coding transcripts were uniquely detected in CSF, accounting for roughly 45% of all transcripts detected in CSF. Approximately 4,377 protein coding transcripts were uniquely detected in plasma. Consistent with these findings, a principal component analysis of biofluid EVs based on the messenger RNA transcriptome revealed that plasma, urine, and CSF EVs cluster separately from each other, highlighting the unique RNA content in these EVs (Figure 2f). To further understand the unique transcriptome content of biofluid derived EVs, we performed biological pathway analysis of differentially expressed genes between biofluids (Supplementary Table 3 & 4). Gene set enrichment analysis (GSEA) of genes upregulated in CSF when compared with plasma revealed significant enrichment of various neurological pathways and dominant contribution from various brain cell types, including neurons, astrocytes, Schwann cells and interneurons (Supplementary Figure 1a). A similar GSEA of genes upregulated in urine when compared to plasma revealed significant enrichment of urological pathways and a dominant contribution originating from various urological cell types (Supplementary Figure 1b). Taken together, these results not only demonstrate the wide diversity of EV transcriptome in biofluids originating from distinct source tissues but also highlight the conservation of RNA species in EVs between biofluids.

### Improved Sequencing Results via Exome & lncRNA Enrichment Using Hybrid-Capture

While WTS presents the diverse landscape of gene expression, we observed that large proportion of reads remain unmappable to human genome, map to non-exonic regions and ERCC spike-ins, pseudogenes and small noncoding RNAs (Figure 2b-d). These result in relatively shallow detection of low-copy genes and transcripts in biofluid EVs, adversely impacting its ability to detect rare biomarkers of interest. Therefore, to focus sequencing reads to regions of interest and further improve our gene detection in biofluid EVs, we employed a whole exome and lncRNA enrichment step on WTS libraries using a hybrid capture-based approach and re-examined the profiles of CSF, plasma, and urine. We observed over 95% of reads mapping to the genome from all biofluids (Figure 3a). Urine samples resulted in over 95% uniquely mapped reads, followed by plasma (75 -85%) and CSF (70 - 80%). This is in sharp contrast to the alignment rates observed with WTS, which demonstrated much lower genome alignment and higher sample-to-sample variability between replicates (Figure 2a). Next, we investigated the genomic origin of reads, which showed remarkable improvement of the proportion of exonic reads from all biofluids compared to WTS. Urine samples, which showed only 45 - 55% of exonic reads in WTS, resulted in the highest proportion of exonic reads (> 90%) among biofluids (Figure 2b, 3b). Plasma EVs which showed between 45 - 55% of exonic reads in WTS, now resulted in ∼90% exonic reads (Figure 2b, 3b). Remarkably, CSF samples, which showed <15% exonic reads in WTS, now resulted in >85% exonic reads (Figure 2b, 3b). In addition, less variation between replicates was also observed for the same biofluid compared to WTS (Figure 2b, 3b). We then evaluated and compared the biotype distribution of exonic reads in our libraries.

**Figure 3.**
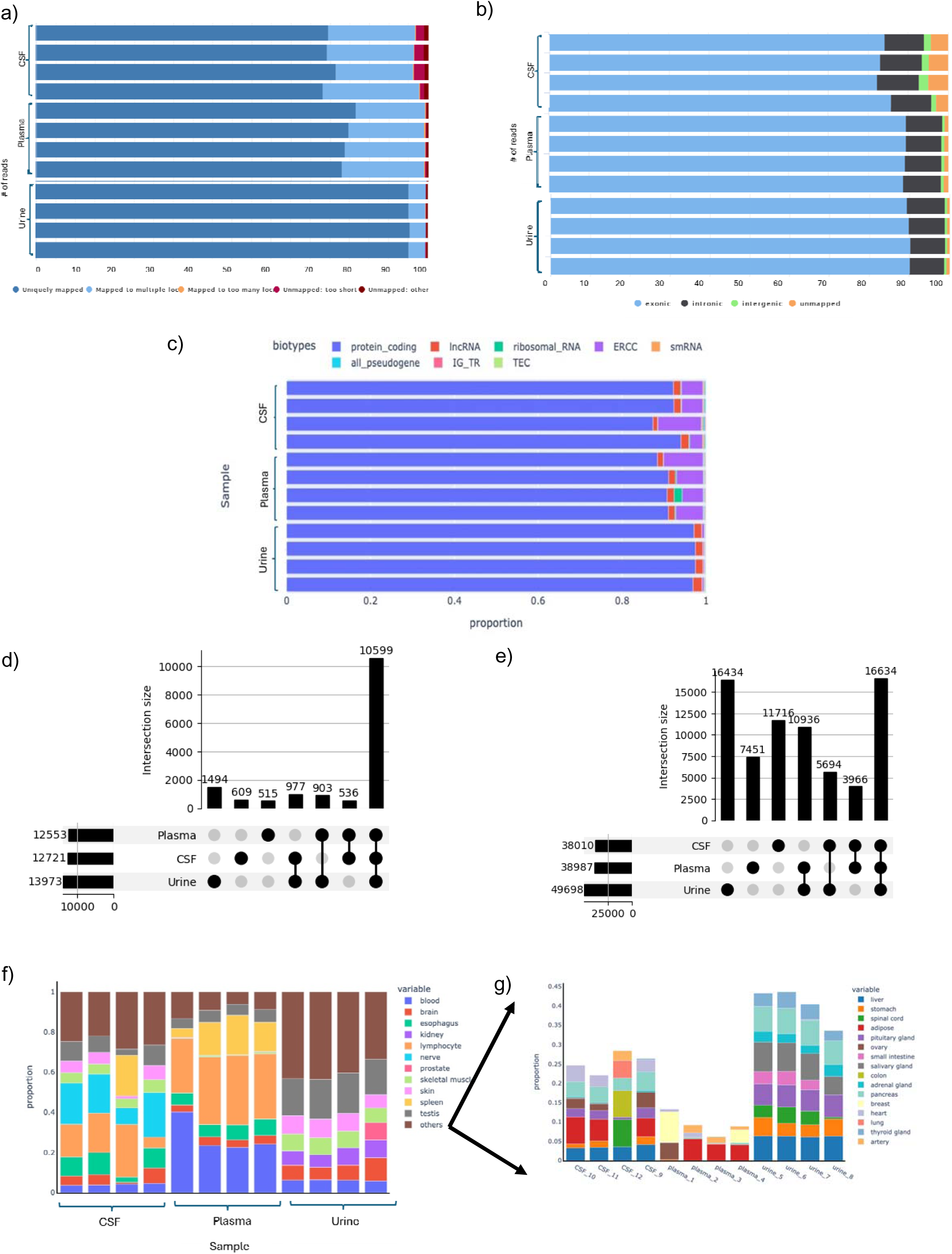
Whole Exome Sequencing across Plasma, Urine, & CSF. a) STAR alignment statistics for plasma, urine, & CSF samples. The x-axis is the percentage of reads. The percentage of uniquely mapped reads varied between biofluids. b) Genomic origin of reads for plasma, urine, & CSF EV samples. The x-axis is the percentage of reads. The percentage of reads mapping to exonic regions is greater than 80% in samples. c) Biotype distributions for plasma, urine, & CSF EV samples. The x-axis is the percentage of reads. The majority of the exonic reads map to protein coding regions in all samples. d) Rate of gene detection across different biofluid derived EVs at a detection threshold of >1TPM. e) Rate of transcript detection across different biofluid derived EVs at a detection threshold of >1 TPM. f) Tissue deconvolution analysis demonstrates the major tissue contribution for each biofluid derived EVs. The y-axis represents proportion of contribution. g) Tissues constituting the “other” category in previous figure.

Approximately 99% of the exonic reads in urine EVs map to protein coding and lncRNA genes, demonstrating further improvement from WTS (∼ 90%) (Figure 2, 3c). Plasma EVs, which showed large sample to sample variability in % of protein coding reads (50 - 85%) in WTS, now resulted in consistent detection of over 85% protein coding reads (Figure 2c, 3c). Similarly, CSF EVs, which showed large sample to sample variability in proportion of protein coding reads (40 - 80%) in WTS, now resulted in consistent detection of over 85% protein coding reads (Figure 2c, 3c). When taken together, these results highlight the excellent on-target rate of enrichment observed by our workflow across different biofluid derived EVs (Supplementary Figure 2).

After demonstrating the marked improvement in focusing of reads to regions of interest using hybrid capture based workflow, we next asked if the increased focusing of reads also resulted in a concomitant improvement in detection of genes and transcripts. As expected, we observed an improved sensitivity of detection of genes from EVs in each biofluid. Plasma EVs, which detected only 9,623 genes in WTS, now detected 12,553 genes post enrichment, demonstrating a 30.44% (additional 2930 genes) improvement in detection of genes (Figure 2d, 3d). Similarly, Urine EVs, which detected 11,977 genes in WTS, now detected 13,973 genes post enrichment, a 16.6% improvement in detection, i.e., additional 1,996 genes (Figure 2d, 3d). In contrast, CSF EVs, which detected 12,532 genes in WTS, only showed an incrementally improved detection of 12,721 genes, an improvement of additional 189 genes only (Figure 2d, 3d). The number of genes detected in all biofluids increased from 7,782 genes in WTS to 10,599 genes in WES, representing a 16% increase in detection, indicating a higher level of overlap between biofluids (Figure 2d, 3d).

Improvement in transcript detection rates from biofluid EVs was found to be even more pronounced than gene detection post-enrichment. For example, plasma EVs, which yielded only 16,552 transcripts in WTS, now produced 38,987 transcripts post-enrichment, demonstrating a 135% improvement in detection, with an additional detection of 22,436 transcripts (Figure 2e, 3e). Similarly, Urine EVs, which yielded 31,395 transcripts in WTS, now produced 49,698 transcripts post enrichment, demonstrating a 53.4% improvement in detection, with an additional detection of 17,303 transcripts (Figure 2e, 3e). In contrast with gene detection improvement, CSF EVs, which yielded 20,080 transcripts in WTS, now produced 38,010 transcripts through WES, showing an 89.29% improvement in detection, resulting in the detection of additional 17,930 transcripts (Figure 2e, 3e). The total number of transcripts detected across biofluids increased from 46,797 to 72,831, a significant jump of 26,034 transcripts (55% higher than the total number of transcripts detected by WTS; Figure 2e, 3e). Similarly, the number of transcripts co-detected in all three biofluids increased from 5,289 transcripts in WTS to 16,634 transcripts post-enrichment, demonstrating a 214.5% increase in detection, with an additional detection 11,345 transcripts (Figure 2e, 3e). Taken together, these results once again highlight the hitherto underappreciated complexity of the EV transcriptome at the transcript level, which seems to be more pronounced than at the gene level. As summarized in Table 1, it establishes the suitability of hybrid capture enrichment in unraveling the complexity of EV transcriptome, both at the gene level and at the transcript level.

**Table 1.** Table comparing RNA-seq QC metrics of WTS & WES for plasma, CSF, and urine. Numbers reported here are the average of four replicates for each biofluid.

|  | Plasma |  | CSF |  | Urine |  |
| --- | --- | --- | --- | --- | --- | --- |
| QC Metrics | WTS | WES | WTS | WES | WTS | WES |
| % Aligned to human genome | 85.6% | 99.4% | 39.1% | 96.9% | 58.1% | 99.5% |
| % Uniquely mapped reads | 22.9% | 79.5% | 11.9% | 74.6% | 21.2% | 95.3% |
| % Exonic reads | 43.5% | 89.3% | 7.7% | 83.8% | 37.3% | 89.9% |
| % Protein-coding of exonic reads | 65.5% | 90.3% | 62.1% | 91.5% | 91.3% | 97.3% |
| % Ribosomal RNA of exonic reads | 5.8% | 0.6% | 1.1% | 0.1% | 0% | 0% |
| Total no. of genes detected | 12,755 | 15,614 | 18,489 | 15,328 | 14,969 | 18,182 |
| No. protein-coding genes detected | 9,623 | 12,553 | 12,532 | 12,721 | 11,977 | 13,973 |
| No. long noncoding RNAs detected | 897 | 1,772 | 2,889 | 1,446 | 1,518 | 3,174 |
| No. protein-coding transcripts detected | 16,552 | 38,987 | 20,088 | 38,010 | 31,395 | 49,698 |

To further explore the tissue of origin of the diverse transcriptome observed in biofluid-derived EVs, we performed a tissue deconvolution analysis of the EV mRNA content of each biofluid, using the Tabula Sapiens database as reference (Figure 3f). As expected, the top three contributors of EV RNA in plasma were blood, lymphocytes and spleen, accounting for over 70% of the RNA content in plasma EVs. Esophagus, testis and brain derived EVs each contributed fewer than 10% of EV RNA. The rest of the tissues in the body accounted for the remaining 5-15% of the RNA content in EVs. In urine, fewer than 5% of RNA was estimated to be of blood-based origin. Testis emerged as the single highest contributor of RNA in urine EVs, while kidney, skin, brain, and skeletal muscles were among the other notable contributors. The rest of the tissues contributed to the remaining 35-45% of the RNA cargo in urine EVs. In contrast, and as expected, nervous system and spleen are the top two contributors of RNA in CSF. Significant contribution was also observed from skeletal muscle, skin, testis and esophagus in CSF. While blood contributed to a small percentage of RNAs in CSF EVs, surprisingly, the contribution of brain derived EVs was also estimated to be < 10%. The rest of the tissues together contributed to the remaining 10 -15% of RNA in CSF EVs.

### Tissue-specific gene panel results in further enrichment while maintaining the fidelity of gene expression

Despite enrichment of protein coding genes and lncRNAs, we observed that the proportion of RNA originating from any solid tissue remained low in biofluid EVs. This is particularly true for plasma EVs, where the hematopoietic contribution accounted for up to 70% of the RNA [58]. The relatively low expression levels of tissue-specific genes and transcripts in plasma EVs therefore makes them hard to detect reliably and poses a major technical challenge for blood based liquid biopsy. We hypothesized that targeted enrichment of tissue specific genes may provide an additional layer of enrichment over WTS and WES, further improving the sensitivity of detection of rare, tissue-specific genes and transcripts. To test this hypothesis, we identified and created bait sets for enrichment of approximately 1,047 genes whose expression levels demonstrated highest levels of enrichment in brain and nervous system, when compared to other tissues in the body, based on the GTEx database [82] (Materials & Methods). To this end, we performed WTS, WES and brain panel enrichment on a set of 20 PD and 20 healthy control plasma samples. Following deduplication based on unique molecular identifiers (UMI) to remove PCR duplicates, we compared the expression level of these genes between WTS and brain panel enrichment. For genes expressed in both WTS and brain panel, we observed an average of 85-fold enrichment of unique molecules of RNA per gene in brain panel with 748 genes (71% of brain panel) exhibiting more than 2-fold enrichment (Figure 4a). Interestingly, not all genes in the panel demonstrated enrichment, with 205 genes showing better detection in WTS. Further investigation revealed that genes showing lack of enrichment had extremely poor transcript coverage and highly fragmented RNA molecules, making enrichment by hybrid capture less efficient (data not shown). Similarly, the comparison of WES and brain panel revealed an enrichment of 23-fold more unique molecules per gene in brain panel, while maintaining excellent correlation of gene expression between them (Figure 4b). While 991 genes were enriched by at least 2-fold in brain panel, there were no genes that were enriched in WES over brain panel.

**Figure 4.**
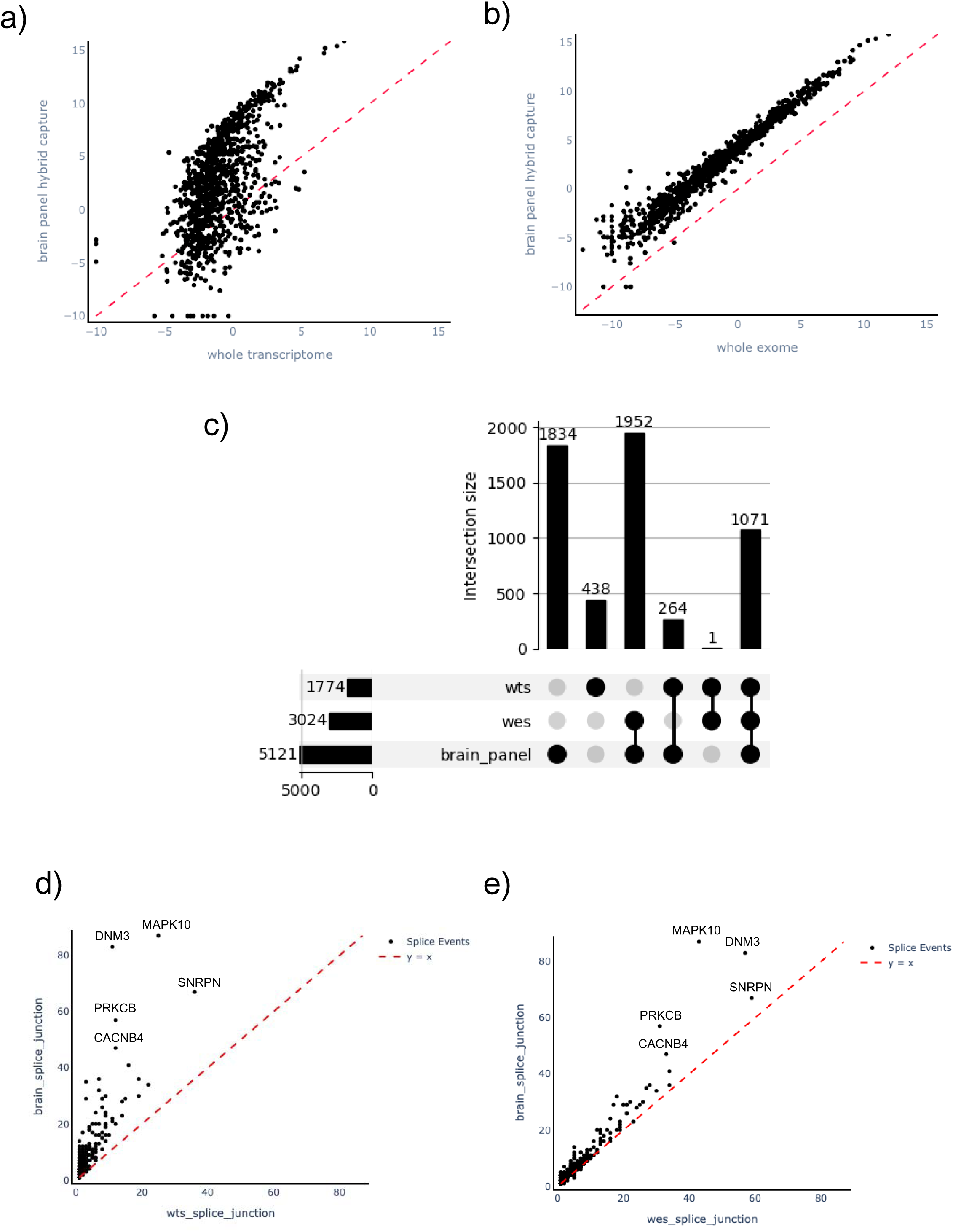
Targeted Brain Panel Sequencing in Parkinson’s Disease & Healthy Plasma Samples. a) Comparison of UMI counts per gene of commonly detected genes in targeted brain panel and WTS. The x-axis is whole transcriptome UMI counts, and the y-axis is brain panel UMI counts. The dashed line marks the identity line. b) Comparison of UMI counts from targeted brain panel sequencing vs WES sequencing for brain panel genes. The x-axis is whole exome UMI counts, and the y-axis is brain panel UMI counts. The dashed line marks the identity line. c) UpSet plot showing the number of transcripts detected in WTS, WES, and brain panel, for all genes that were detected across the three platforms. d) Scatter plot showing the UMI counts per splice event of genes detected in both brain panel and WTS. The x-axis is whole transcriptome UMI counts, and the y-axis is brain panel UMI counts. The dashed line marks the identity line. e) Scatter plot showing the UMI counts per splice event of genes detected in both brain panel and WES. The x-axis is whole exome UMI counts, and the y-axis is brain panel UMI counts. The dashed line marks the identity line.

Next, we compared the detection of transcripts across the three platforms, limiting ourselves only to the 992 genes commonly detected across platforms. We observed that the brain panel detected more transcripts than any other platform with 5,121 transcripts, resulting in 188.7% increase (additional 3,347 transcripts) in detection of transcripts from WTS (1,774 transcripts) and 69.3% increase in detection (additional 2,097 transcripts) of transcripts when compared to WES (3024 transcripts) (Figure 4c). Finally, we analyzed the no. of splice events detected per gene across the three platforms. Our analysis revealed that brain panel detected on avg. 3.1-fold more splicing events per gene than WTS (Figure 4d) and 1.4-fold more splicing events after WES (Figure 4e). Taken together, these observations point to the tremendous improvement & highly sensitive detection of rare, tissue specific genes, transcripts & splice junctions achieved by targeted brain panel, when compared with WTS & WES. An IGV plot of MAPK10, one of the genes that showed major enrichment in splice junction detection, exemplifies these improvements described above in Supplementary Figure 3.

### Performance of WTS, WES and Brain panel gene expression on Parkinson’s disease biomarker discovery

We tested WTS, WES and brain panel for their performance and suitability for biomarker discovery of Parkinson’s disease (PD) from a set of 20 PD and 20 healthy control plasma EV samples. Parkinson’s disease is a progressive neurological disorder that affects movement, leading to characteristic motor symptoms like tremors, slowness of movement, stiffness, and postural instability [83, 84]. While no specific test definitively diagnoses Parkinson’s, Parkinson’s disease diagnosis is primarily based on a clinical evaluation, including a medical history and neurological examination by a neurologist [83, 85]. Differential expression analysis between PD and healthy controls identified 5 significantly differentially expressed (DEx) genes from the brain panel, 9 DEx genes from WES and 653 DEx genes from WTS (Figure 5a). While none of the DEx genes were shared across the three platforms, one DEx gene was shared across any two platforms (Figure 5a). For example, POU3F2 gene was shared between WES & brain panel, the gene STUM was shared across brain panel & WTS, while the gene HS3ST4 was shared between WTS and WES. Approximately 60% of the 653 DEx genes that were identified in WTS were protein coding genes, with lncRNAs, pseudogenes, small RNA, TEC and Misc RNAs constitute the remaining ∼40% of the DEx genes between PD and healthy (Figure 5b). As expected, WES DEx genes consisted of both protein coding genes and lncRNAs, while brain panel DEx genes were entirely mRNAs (Figure 5b). Feature selection analysis on WTS DEx genes using Boruta identified six genes with maximal contribution to separation between PD and healthy (Table 2), consisting of five protein coding genes and one pseudogene. The performance of a classifier using these features to separate PD from healthy using Naïve Bayes model resulted in a leave-one-out-cross-validation (LOOCV) AUC of 0.93 (Figure 5c & d). Similarly, feature selection on WES DEx genes identified three protein coding genes, namely GPR146, IL17D & MYOM3 (Table 2), resulting in a classifier performance AUC of 0.88 (Figure 5c & e). Feature selection identified four protein coding genes from brain panel DEx genes, namely POU3F2, PPP2R2B, SPTBN2 and STUM (Table 2), with a classifier performance AUC of 0.87 (Figure 5c & f).

**Figure 5.**
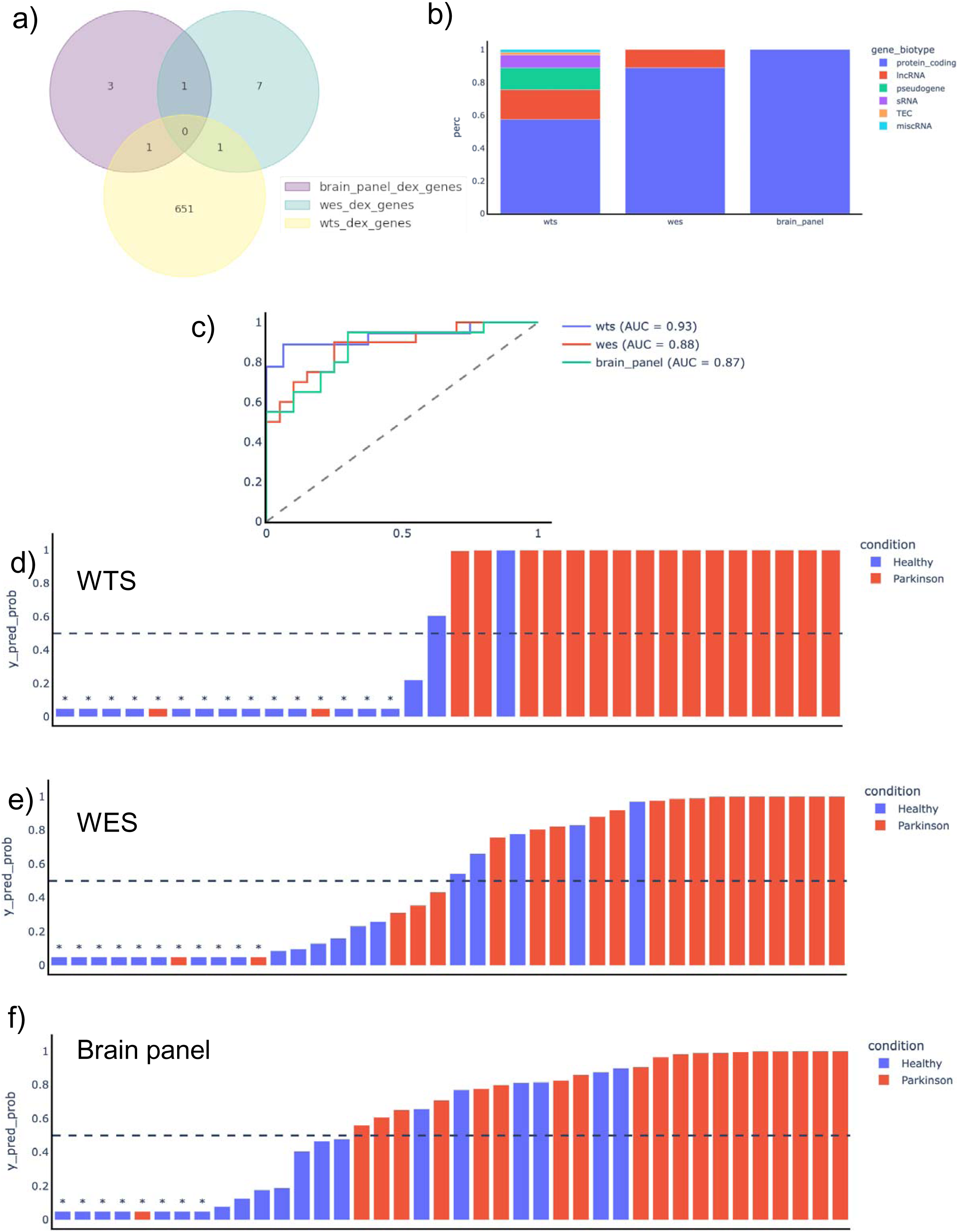
Biomarker Discovery of Parkinson’s Disease and Healthy Plasma Samples using Differential Gene Expression. a) Overlap of differentially expressed genes between Parkinson’s and healthy plasma EVs profiled using WTS, WES, and targeted brain panel sequencing. No Dex genes were identified in all three platforms. b) Biotype distribution for differentially expressed genes by WTS, WES, and brain panel sequencing. The y-axis is percentage of reads. c) ROC curves of Parkinson’s gene signatures from WTS, WES, and brain panel showing AUC values of 0.84, 0.84, and 0.85 respectively. d) Waterfall plot demonstrating the performance of Parkinson’s gene expression-based signature using WTS. Healthy samples are in blue, while Parkinson’s samples are in orange. Dotted line represents cutoffs to achieve sensitivity and specificity indicated in figure. Samples with an asterisk indicate sample with predicted prob<0.05 and were imputed as 0.05 to be visible in the plot. e) Waterfall plot demonstrating the performance of Parkinson’s gene expression-based signature using WES. Healthy samples are in blue, while Parkinson’s samples are in orange. Dotted line represents cutoffs to achieve sensitivity and specificity indicated in figure. Samples with an asterisk indicate sample with predicted prob<0.05 and were imputed as 0.05 to be visible in the plot. f) Waterfall plot demonstrating the performance of Parkinson’s gene expression-based signature using brain panel sequencing. Healthy samples are in blue, while Parkinson’s samples are in orange. Dotted line represents cutoffs to achieve sensitivity and specificity indicated in figure. Samples with an asterisk indicate sample with predicted prob<0.05 and were imputed as 0.05 to be visible in the plot.

**Table 2.** Table listing the differentially expressed genes selected for the classifier for WTS, WES, and brain panel sequencing.

| WTS (k=6) | WES (k=3) | Brain panel (k=4) |
| --- | --- | --- |
| 'DNAH5', 'ENSG00000224629', 'FIG4', 'KCNK16', 'SAPCD2', 'TMEM185B' | 'GPR146', 'IL17D', 'MYOM3' | 'POU3F2', 'PPP2R2B', 'SPTBN2', 'STUM' |

### Performance of WTS, WES and Brain panel splice variants on Parkinson’s disease biomarker discovery

Next, we tested WTS, WES and brain panel splice variants for their performance and suitability for biomarker discovery from the same set of 20 PD and 20 healthy control plasma EV samples. Differential splice variants analysis identified 69 splicing events from the brain panel, 1552 splicing events from WES and 201 splicing events from WTS with significantly different percent spliced in PSI scores (Figure 6a). Interestingly, very little overlap of significantly different splicing events between PD and healthy controls was observed between the three methods, with only one splicing event shared across the three platforms (CAMKK2_SE_JC), 26 common between brain panel and WES, and 47 common between WES and WTS (Figure 6a). Unlike gene expression in WTS, WES platform resulted in the detection of the largest no. of significantly differential splicing events (Figure 6a, Figure 5a). Feature selection analysis on differential splicing events using Boruta identified 6 events with maximal contribution to separation between PD and healthy from each of the platforms (Table 3, Supplementary Figure 4). The performance of a Naïve Bayes classifier using these features to separate PD from healthy result in LOOCV AUC of 0.92, 0.90 and 0.91 for WTS, WES and brain panel, respectively (Figure 6b-e).

**Figure 6.**
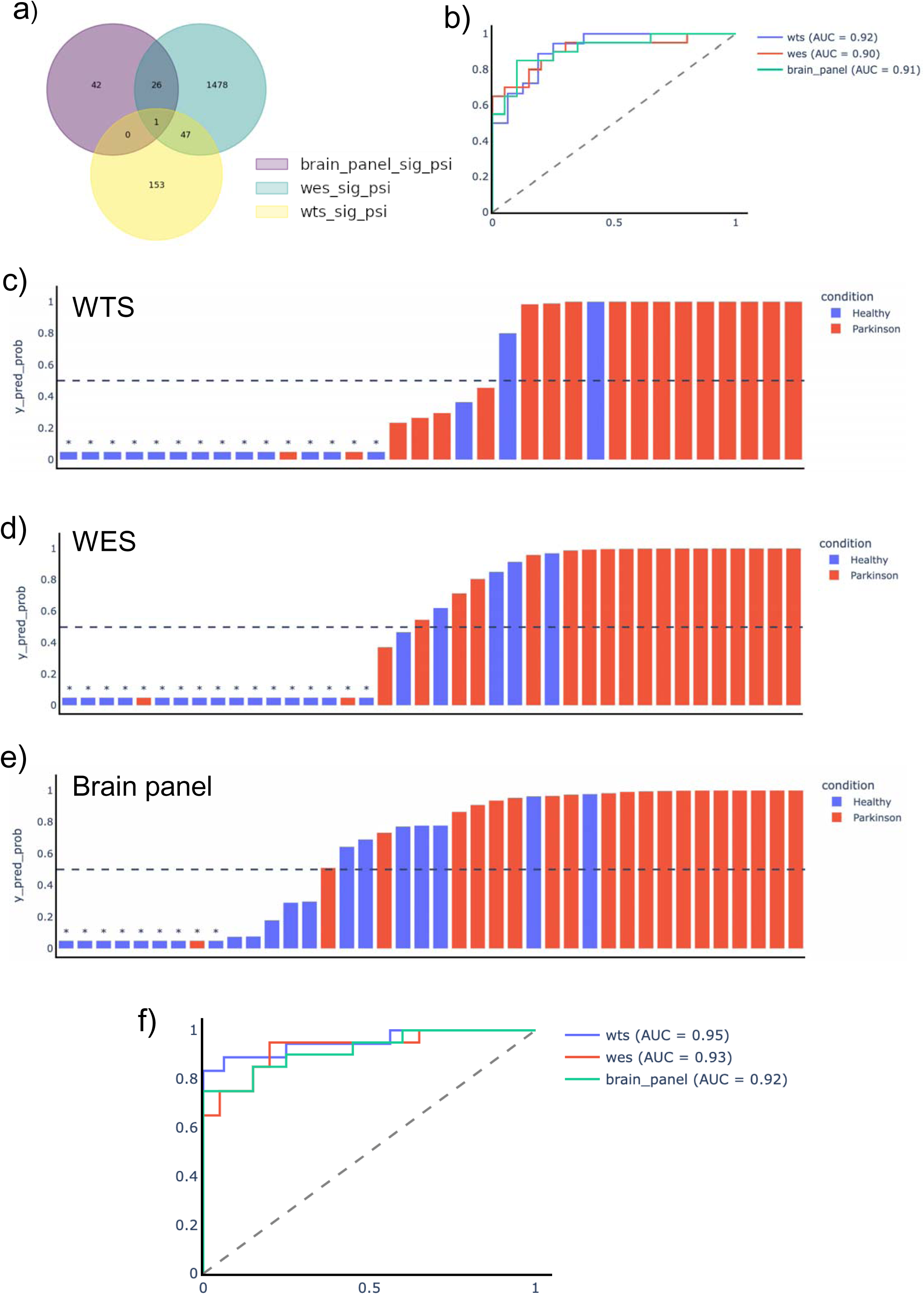
Biomarker Discovery of Parkinson’s Disease and Healthy Plasma Samples using Splice Event Analysis and Multiomic Performance. a) Overlap of differential splice events detected using WTS, WES, and targeted brain panel sequencing between Parkinson’s Disease and healthy plasma samples. One splicing event was shared across the three platforms. b) ROC curves of Parkinson’s splice event signatures from WTS, WES, and brain panel showing AUC values of 0.92, 0.90, and 0.91 respectively. c) Waterfall plot demonstrating the performance of Parkinson’s splice event-based signature for differential splice events in WTS. Healthy samples are in blue, while Parkinson’s samples are in orange. Dotted line represents cutoffs to achieve sensitivity and specificity indicated in figure. Samples with an asterisk indicate sample with predicted prob<0.05 and were imputed as 0.05 to be visible in the plot. d) Waterfall plot demonstrating the performance of Parkinson’s splice event-based signature for differential splice events in WES. Healthy samples are in blue, while Parkinson’s samples are in orange. Dotted line represents cutoffs to achieve sensitivity and specificity indicated in figure. Samples with an asterisk indicate sample with predicted prob<0.05 and were imputed as 0.05 to be visible in the plot. e) Waterfall plot demonstrating the performance of Parkinson’s splice event-based signature differential splice events genes in brain panel sequencing. Healthy samples are in blue, while Parkinson’s samples are in orange. Dotted line represents cutoffs to achieve sensitivity and specificity indicated in figure. Samples with an asterisk indicate sample with predicted prob<0.05 and were imputed as 0.05 to be visible in the plot. f) Multiomic signatures obtained from combining gene expression and splice event features from WTS, WES, and brain panel, respectively demonstrated AUC values of 0.95, 0.93, and 0.92.

**Table 3.** Table listing the differential splice events selected for the classifier for WTS, WES, and brain panel sequencing.

| WTS (k=6) | WES (k=6) | Brain panel (k=6) |
| --- | --- | --- |
| MAP3K7CL_MXE_JC_1172',<br>'FLII_RI_JC_4232',<br>'CIRBP_RI_JC_5296',<br>'BSG_RI_JC_7998',<br>'PKIG_SE_JC_25145',<br>'SEPTIN2_SE_JC_45791 | MCUR1_MXE_JC_2311',<br>'ZNF592_MXE_JC_8945',<br>'DEK_MXE_JC_10941',<br>'UGP2_SE_JC_39179',<br>'ACTN1_SE_JC_46748',<br>'DCTN4_SE_JC_58567 | PEA15_MXE_JC_2118',<br>'PPP3CA_SE_JC_17079',<br>'SNCA_SE_JC_24885',<br>'DNM3_SE_JC_40598',<br>'IPCEF1_SE_JC_42822',<br>'DMTN_SE_JC_52642' |

Finally, we investigated if there is complementarity of signal between gene expression and splice variants-based biomarkers and if further improvement in the diagnostic performance of the models can be achieved by combining the top biomarkers of gene expression and splice variants. Boruta feature selection from the top feature selected genes and splice variants resulted in the identification of a multiomic feature set combining top expression and splice variant markers for each platform (Table 4). The performance of a classifier using these multiomic features to classify PD from healthy using LOOCV resulted in a marginal improvement of AUC to 0.95, 0.93 and 0.92, respectively (Figure 6f).

**Table 4.** Table listing the multiomic features selected for the classifier for WTS, WES, and brain panel sequencing selected from the differentially expressed genes and differential splice events used in their respective classifiers.

| WTS (k=6) | WES (k=6) | Brain panel (k=6) |
| --- | --- | --- |
| 'PKIG_SE_JC_25145', 'DNAH5',<br>'FIG4', 'KCNK16', 'SAPCD2',<br>'TMEM185B' | 'MCUR1_MXE_JC_2311',<br>'ZNF592_MXE_JC_8945',<br>'DEK_MXE_JC_10941',<br>'UGP2_SE_JC_39179', 'GPR146',<br>'MYOM3' | 'PEA15_MXE_JC_2118',<br>'PPP2R2B',<br>'SNCA_SE_JC_24885',<br>'DNM3_SE_JC_40598',<br>'IPCEF1_SE_JC_42822',<br>'SPTBN2' |

## Discussion

In this study, we performed a comprehensive profiling and comparative analysis of EV long RNAs by WTS in three biofluids: plasma, urine and CSF. The type of biological information we obtained per sample differed vastly by biofluid, including the quantity and quality of exoRNA they contain, and analysis of WTS revealed trends unique to each body fluid studied.

Specifically, we observed a large proportion of unmapped reads and low exoRNA content in CSF compared to other body fluids. These characteristics may be due to the pervasive RNA fragmentation inherent to CSF and be responsible for the enriched ERCC reads in CSF samples compared to other biofluids. In contrast, the high quality and less fragmented exoRNA in urine is thought to contribute to the higher protein-coding mapping and increased rRNA depletion compared to other fluids. A combined total of approximately 15,000 unique genes representing ∼25% of all known human genes were detected in any biofluid, with over 7700 genes shared among the three biofluids. Similarly, a total of over 46,000 transcripts, representing ∼28% of all known human transcripts, were detected in any biofluid, with only 5289 transcripts shared across all three biofluids. This is interesting, considering over 50% of genes detected are shared across biofluids, while only 10% of transcripts are shared across biofluids. Consistent with observations in cellular and tissue level transcriptomes [86], these results reveal that the diversity of transcriptome in biofluid EVs is higher at the transcript level than at the gene level, underscoring a hidden layer of complexity of EV transcriptomes that has not yet been appreciated fully. It further suggests some universality in the way exoRNA cargo is sorted into EVs and distributed to the specific biofluids. Large number of genes and transcripts were also found to be uniquely detectable in each biofluid, with CSF showing the highest number of unique genes while urine showed the highest number of unique transcripts. Gene set enrichment and pathway analysis all pointed to differential contribution of EVs from source tissues in these biofluids as the underlying cause of this diversity. Therefore, when selecting a biofluid for liquid biopsy or biomarker discovery, proximity of the tissue to the biofluid should be taken into consideration.

Taken together, these results underscore the wide diversity of EV transcriptome in biofluids originating from distinct source tissues and at the same time also highlight the conservation of RNA species in EVs between biofluids.

The large proportion of reads unmappable to human genome, reads mapping to non-exonic regions and ERCC spike-ins, pseudogenes and small noncoding RNAs in WTS adversely affect its sensitivity and pose a challenge to detect rare biomarkers of interest in biofluid EVs. To address these challenges, we described a workflow that utilizes enrichment of all protein coding genes and lncRNA using hybrid capture on WTS libraries. Our results demonstrated remarkable focusing of reads to regions of interest using hybrid capture, leading to improved RNASeq QC metrics, reduced sample to sample variability and highly consistent on-target rate of enrichment. These improvements resulted in a concomitant improvement in genes and transcript detection from biofluid EVs, with improvement in transcript detection was found to be much more pronounced than gene detection post enrichment. Moreover, to demonstrate the robustness of this workflow, we performed exhaustive reproducibility studies by profiling 72 biological replicates of plasma EV samples by four independent operators (Supplementary Figure 5). In this study, we observed excellent transcriptome wide correlation of gene expression (> 0.95, Pearson’s coefficient), correlation of ERCC spike-ins levels (> 0.97, Pearson’s), highly consistent mapping statistics and biotype distribution, resulting in consistent detection of over 13,000 genes and 32,000 transcripts in almost all replicates. Importantly, over 3,200 genes and over 7,000 transcripts were consistently detected with high coverage (> 80% exonic length coverage) in almost all replicates, representing ∼20-25% of total detected genes and transcripts. These results contrast previous reports of enrichment of 3’UTR fragments in EVs and underscore a major improvement achieved by this method, enabling splice variant analysis of EV RNA [87].

Tissue deconvolution analysis pointed to differential contribution of tissue EVs in different biofluids; however, caution should be exercised while interpreting these results. Deconvolution analysis relies on publicly available tissue single cell RNASeq databases which have their own limitations. Such analysis assumes RNA distribution in tissues is reliably maintained in biofluids and ignores the possibility of specific RNA sorting in EVs. It also assumes that each tissue has the same ability of releasing EVs in all biofluids and ignores proximity or barriers of some tissues to EV release in specific biofluids (e.g., blood-brain barrier). Taken together, these results once again highlight the hitherto underappreciated complexity of the EV transcriptome at the transcript level. Furthermore, it establishes the suitability of hybrid capture enrichment in unraveling the complexity of EV transcriptome, both at the gene and at the transcript level.

Tissue-specific enrichment and profiling of EVs from plasma is complicated by the abundance of the hematopoietic EVs in plasma and represents one of the holy grails of liquid biopsy. Majority of the efforts have focused on enrichment of tissue specific EVs from biofluids using antibodies against tissue specific surface proteins on EV membranes. However, this approach has yielded limited success to date and on some occasions has been controversial[60, 61]. Challenges include identification and detection of true tissue-specific surface markers on EVs and distinguishing them from cell-free proteins, availability of antibodies, rarity of these EVs in plasma, lack of sensitive techniques for RNA profiling post enrichment, among others. In this study, we presented an alternate approach for tissue specific RNA profiling that relies on prior identification of these genes using publicly available tissue RNASeq databases such as GTEx & Tabula Sapiens. The brain panel results presented in this study is meant to be a proof of concept yet clearly demonstrate the tremendous enrichment of brain derived genes, transcripts and splicing events that can be achieved through this approach. This approach is also easily generalizable to other tissues and represents one of the most sensitive ways of profiling rare tissue-specific transcripts from plasma. However, such an approach is not without its own limitations. The need for prior identification of tissue specific genes relies on publicly available tissue RNASeq databases and limits its throughput for unbiased biomarker discovery. Moreover, most genes are not expressed in a tissue specific manner, therefore, biomarkers that do not meet this criterion might be missed through this approach. Significant number of brain panel genes were also found to be more abundant in WTS than brain panel (Figure 4a). However, these transcripts were most likely highly fragmented in nature, leading to failure of hybrid capture enrichment.

Next, we tested the suitability and performance of all three RNASeq strategies (WTS, WES and brain panel) for biomarker discovery in liquid biopsy, using Parkinson’s disease as a case study. Biomarker discovery of PD using plasma EVs, a neurodegenerative disease of the CNS, presents a particularly challenging case study as the blood brain barrier makes brain derived EVs an extreme rarity. Yet, when looking at gene expression biomarkers, all three RNA profiling approaches resulted in similar performance, as DeLong’s test shows no significant difference between AUCs (data not shown). Interestingly, very little overlap in DEx genes was observed between the three methods, demonstrating the different focus of each approach. In WTS, while literature review did not point to any known relevance of five out of six feature selected genes, interestingly, FIG4 has been associated with a rare form of Parkinsonism that often co-occurs with Charcot-Marie-Tooth disease type 4J (CMT4J) [88, 89]. Similarly, an exhaustive literature survey of feature selected genes in WES showed limited relevance in PD pathophysiology. While MYOM3 & IL17D has no known relevance in Parkinson’s disease, other members of the IL17 cytokine family, specifically IL17A, has been linked to neuroinflammation, which is a key feature of PD [90, 91]. Similarly, GPR146 has been suggested to be a therapeutic target for PD [92, 93]. However, in contrast to WTS and WES, in-depth literature review of feature selected genes from brain panel revealed many of these genes have been suggested to play a role in PD pathophysiology [94–101], other neurodegenerative diseases like Alzheimer’s and Spinocerebellar ataxia [97, 102–104], mechanical sensing in proprioceptive neurons & neuroinflammation [94].

In WTS, Pathway analysis of DEx genes between PD and healthy did not point to any pathway related to Parkinson’s disease. While GO analysis of genes upregulated in PD pointed to pathways related to Globins and oxygen transport, pathway analysis of down regulated genes pointed to a heterogenous mix of pathways, including chemotaxis, cell adhesion, matrix metallopeptidase secretion and endothelial cells (Supplementary Figure 6a, 6b). In WES, GO analysis of upregulated genes did not point to any relevant pathway (Supplementary figure 7a), while GO analysis of downregulated genes did point to a subset of pathways related to nervous system, including, synapse structure, assembly and organization, astrocytes, hypothalamus cell differentiation, neuronal fate specification and neurotrophins binding, among others (Supplementary figure 7b). In contrast to WTS and WES, in brain panel, GO analysis of genes upregulated in PD pointed to tremendous enrichment of several pathways related to nervous system, including Purkinje cells synapses, morphogenesis & development, cerebellar cortex morphogenesis and development, synapse & excitatory neurons, among others (Supplementary figure 8a). It is worth noting that Purkinje cells and cerebellum have been heavily implicated in Parkinson’s disease and other movement related disorders [105–107]. In genes downregulated in PD, GO analysis revealed tremendous enrichment of pathways related to mechanosensory behavior, hypothalamus cell differentiation and development, excitatory neurons, synapse assembly, maturation and transmission, and several other pathways relevant to CNS (Supplementary figure 8b). It is worth noting that while some of these pathways were also observed in WES, the level of enrichment is significantly higher in brain panel compared with WES. Taken together, and consistent with our observations on feature selected genes described above, pathway analysis of dysregulated genes in PD showed highest enrichment of CNS and PD related pathways in brain panel, followed by WES which shows some brain relevance and finally by WTS which showed least relevance to CNS and PD.

The use of splice variants-based biomarker discovery in liquid biopsy for a neurodegenerative disease like PD represents a completely novel feature of this study. Like gene expression, very little overlap of differential splicing events was observed across the three platforms (figure 6a). Like gene expression, in-depth literature review of the feature selected events in WTS and WES revealed the splicing events to be novel, with no known relevance in Parkinson’s disease. In contrast, feature selected splicing events from brain panel were also novel, however, the genes themselves in which they occurred were found to be most relevant to PD pathophysiology[108–119]. For example, SNCA, which encodes the protein alpha synuclein which is a major component of Lewy bodies, has been strongly associated with PD and is perhaps the best known and most studied biomarker of PD (supplementary figure 9) [116–119]. The gene PEA15 has been linked to PD through its role in insulin resistance and its potential to improve dopaminergic pathways in the brain [108, 109]. Specific mutations in the genes PPP3CA, DNM3 and DMTN has been previously linked with Parkinson’s, although the exact mechanisms are still being explored [110–112]. Taken together, these results highlight that even if each of the EV RNASeq modalities, i.e., WTS, WES and brain panel resulted in a comparable performance, the underlying genes and pathways from the brain panel were most relevant to CNS and disease pathophysiology, maximizing the confidence on successful scrutiny of these biomarkers in larger sample cohort. In contrast, the signatures identified by WTS & WES are less likely to pass scrutiny in a larger cohort, although it remains to be empirically established in a larger cohort.

It is worth noting that across each platform, the performance of the models based on splice variants outperformed the models based on gene expression. This is significant, as splice variants-based biomarkers have been explored to a limited extent in liquid biopsy, not just in PD, but across other disease conditions as well. Therefore, the use of splice variants as a platform for liquid biopsy is currently underappreciated and may open a treasure trove of novel biomarkers across various conditions in future [120].

Combining gene expression and splice variant biomarkers to obtain a multiomic signature that exploits potential signal complementarity and obtain improved performance is yet another novelty of this study. While our results demonstrated only a nominal improvement in AUC, the small sample numbers in this study heavily limited our ability to exploit this approach fully.

While this study serves as a proof-of-concept of this approach, future studies with larger sample sizes will be needed to explore the power and potential of multiomics to its full extent.

## Materials and Methods

### Sample Details

Whole transcriptome sequencing (WTS) and whole exome RNA sequencing (WES) was performed on four pools of plasma, urine, and CSF from healthy individuals. Each biofluid pool is comprised of ten healthy individuals. Samples were stored at -80C prior to further processing. Samples were filtered by 0.8 µm filter prior to EV isolation. WTS, WES, and brain panel sequencing was also performed on 20 Parkinson’s disease patients and 20 healthy control plasma samples, which were age and gender matched, and procured from a commercial biobank (Precision for Medicine) (Supplementary Table 1).

### RNA Isolation

Total RNA was purified from 2 mL of filtered plasma or CSF using Exosome Diagnostic’s proprietary ExoLution workflow [121]. Total RNA was purified from 10 mL of filtered urine using Exosome Diagnostic’s proprietary UPrep workflow [122]. RNA was assessed for quality using Agilent Bioanalyzer with the RNA Pico 6000 Kit (Agilent Technologies). Isolated RNA was stored at -80 °C prior to library generation.

### Library Construction

#### Whole Transcriptome Sequencing (WTS)

Library construction for profiling EV long RNA was performed using Exosome Diagnostics’ proprietary EV long RNA sequencing workflow. Briefly, isolated RNA from Plasma, CSF, or Urine as described above) was treated with DNase to remove any co-purified DNA present in the sample. Synthetic RNA spike-in controls (ERCC spike-in, cat no. 4456740, Thermo Fisher Scientific,) were then added to each sample. Total RNA was then reverse transcribed using a combination of random hexamers and oligo dT primers. Adapter addition was performed using a PCR-based approach. Libraries were purified and size selected via two rounds of AMPureXP® beads (cat no. A63881, Beckman Coulter). Libraries were ribodepleted to remove ribosomal cDNA. The final ribodepleted libraries were then amplified by PCR, followed by clean up using AMPureXP® beads. Libraries were quantified using the Qubit 1X dsDNA HS Assay Kit (cat no. Q33231, Thermo Fisher Scientific) and the BioAnalyzer HS-DNA Chip (cat no. 5067-4626, Agilent Technologies). Final libraries were pooled to equimolar concentrations and sequenced on Illumina® NextSeq500 using 2 × 151 cycles read length chemistry with PhiX control, achieving an average sequencing depth of 40M reads.

#### Whole Exome RNA Sequencing (WES)

WES libraries for plasma, urine, & CSF were constructed as described above with the only modification being no ribodepletion was performed in these libraries. Libraries were quantified using the Qubit 1X dsDNA HS Assay Kit (Thermo Fisher Scientific) and the BioAnalyzer HS-DNA Chip (Agilent Technologies). Eight libraries were pooled by equal mass to a total yield of 1500 ng. Pooled libraries were then hybridized to custom bait sets against all protein-coding genes with UTRs, long noncoding RNAs, and ERCCs using Agilent Technologies’ proprietary probe design software. Hybridized libraries were captured and washed as per Agilent’s workflow. Final libraries were then amplified using PCR and cleaned using SPRI beads.

Captured libraries were quantified using the Qubit 1X dsDNA HS Assay Kit (Thermo Fisher Scientific) and the BioAnalyzer HS-DNA Chip (Agilent Technologies). Final libraries were pooled to equimolar concentrations and sequenced on Illumina® NextSeq500 using 2 × 15<u>1</u> cycles read length chemistry with PhiX control, achieving an average sequencing depth of 30M reads.

#### Targeted Brain Panel Sequencing

Total RNA sequencing libraries were prepared as described above. Hybrid capture based enrichment was performed using brain panel bait sets as described for WES above. For the brain panel, the top 1047 protein-coding genes with the highest enrichment (Tmax % > 50 %) in the brain and nervous system and a minimum TPM of 100 were identified based on GTEx database.

Tmax % is calculated with the following equation:

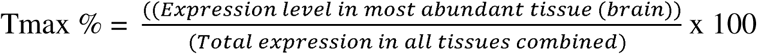

Bait sets were designed using Agilent’s proprietary probe design software. Final libraries were pooled to equimolar concentrations and sequenced on Illumina® NextSeq550 using 2 × 151 cycles read length chemistry with PhiX control, achieving an average sequencing depth of 40M reads.

### Bioinformatic Analysis

#### Data processing

Gene-level expression data was obtained using our proprietary RNASeq pipeline, which is designed for EV RNA sequencing. STAR[123] and salmon[124] were used for sequencing alignment and quantification in the pipeline. PCR duplicates were removed using UMI-tools[125] to obtain unique molecular counts in the sample.

#### Differential expression analysis and pathway analysis

Gene expression data was filtered to remove genes with no read count across all samples. Differential expression analysis was performed using DESeq2[126] and differential expression genes (DEGs) were selected with BH-adjusted p-value of 0.05. Gene set enrichment analysis (GSEA) was then performed using DESeq2 results with log2 fold change corrected by approximate posterior estimation for GLM (apeglm)[127] to remove noise from real gene expression differences. Gene ontology database from MSigDB [128, 129]was used for pathway annotation.

#### Differential splice event analysis

Splice variant analysis was conducted at both the transcript level and splice event level using a custom Nextflow pipeline. Transcript quantification derived from Salmon (v1.9.0) using the previously mentioned gene expression pipeline were used to calculate Percent Spliced In (PSI) values of transcript using SUPPA [130] (v2.3). Aligned reads generated from STAR (v2.7.10a) using the previously mentioned gene expression pipeline were provided as input to RMATs [131](v4.1.2) to detect splice events to quantify PSI values of splice events belonging to one of the five following categories: skipped exon (SE), alternative 5’ splice sites (A5SS), alternative 3’ splice sites (A3SS), mutually exclusive exons (MXE), or retained intron (RI).

#### Feature selection and model fitting

Differential expression genes (DEGs) obtained from DESeq2 were initially utilized as the feature set to create a signature and build a model for distinguishing Parkinson’s disease samples from healthy samples. Subsequently, the Boruta algorithm was employed on these DEGs to pinpoint a subset of genes that hold the utmost relevance for classification purposes.

A nested cross-validation (CV) approach was adopted to finalize the feature selection process in the inner CV and to fit a Gaussian Naïve Bayes model. This involved restricting the maximum number of features in the model to 6. Area Under the Curve (AUC) was computed from leave-one-out cross-validated predictions in the outer CV.

## Code Availability

https://github.com/exosomedx/biofluid_ev_reveal.git

