## Supplemental Figures for "Deep Profiling of EV Long RNAs Reveals Biofluid-Specific Transcriptomes and Splicing Landscapes"

### Slide 1
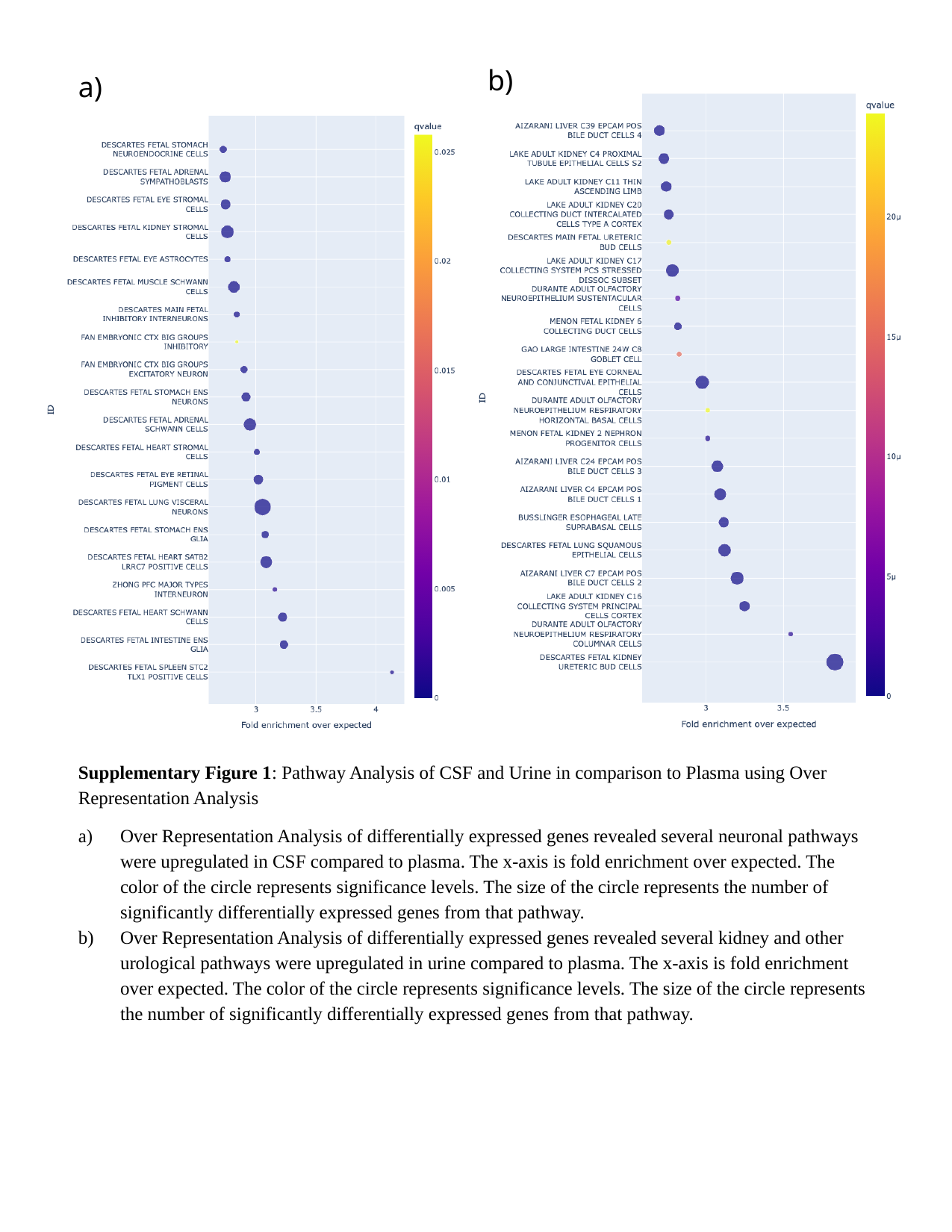

b)
a)
Supplementary Figure 1: Pathway Analysis of CSF and Urine in comparison to Plasma using Over Representation Analysis
Over Representation Analysis of differentially expressed genes revealed several neuronal pathways were upregulated in CSF compared to plasma. The x-axis is fold enrichment over expected. The color of the circle represents significance levels. The size of the circle represents the number of significantly differentially expressed genes from that pathway.
Over Representation Analysis of differentially expressed genes revealed several kidney and other urological pathways were upregulated in urine compared to plasma. The x-axis is fold enrichment over expected. The color of the circle represents significance levels. The size of the circle represents the number of significantly differentially expressed genes from that pathway.

### Slide 2
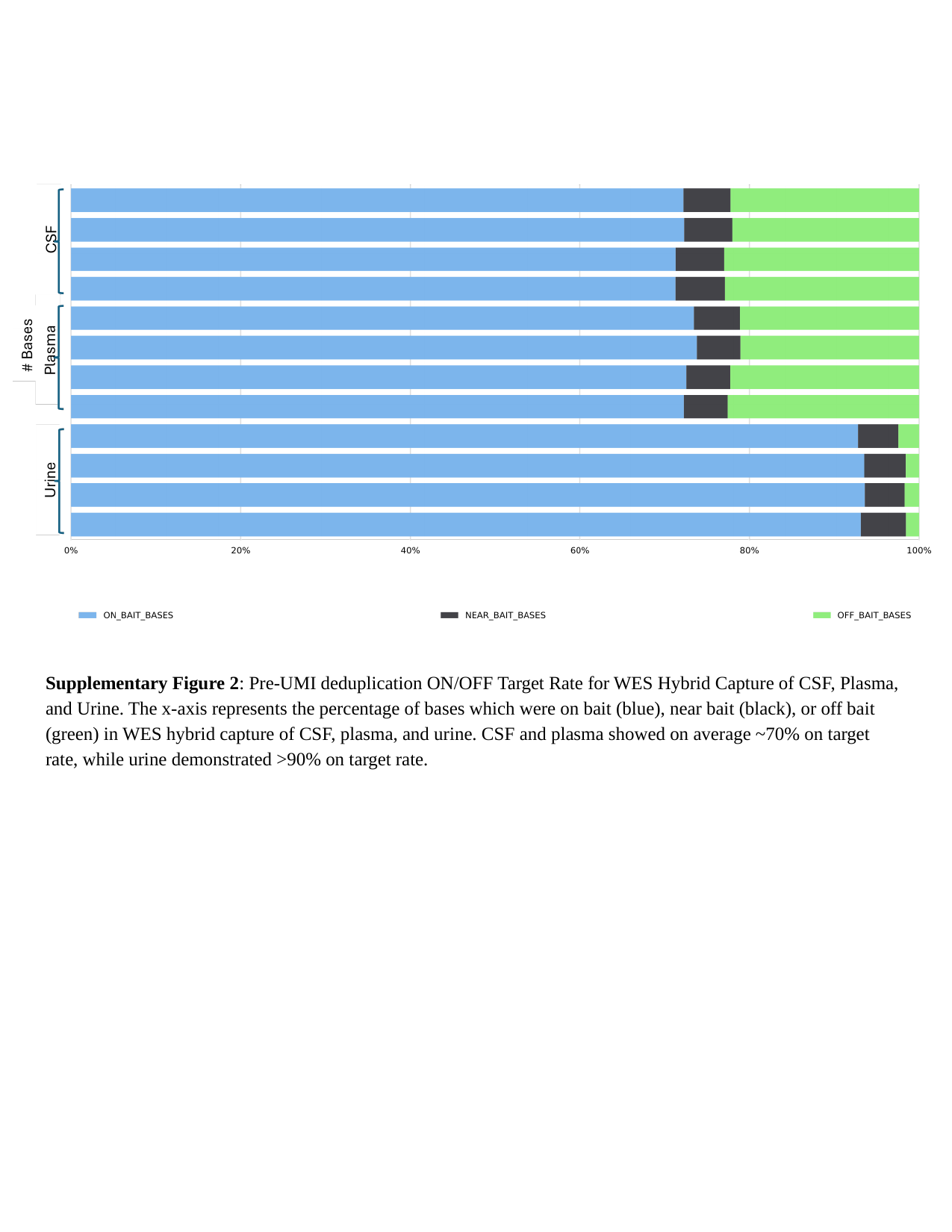

Supplementary Figure 2: Pre-UMI deduplication ON/OFF Target Rate for WES Hybrid Capture of CSF, Plasma, and Urine. The x-axis represents the percentage of bases which were on bait (blue), near bait (black), or off bait (green) in WES hybrid capture of CSF, plasma, and urine. CSF and plasma showed on average ~70% on target rate, while urine demonstrated >90% on target rate.

### Slide 3
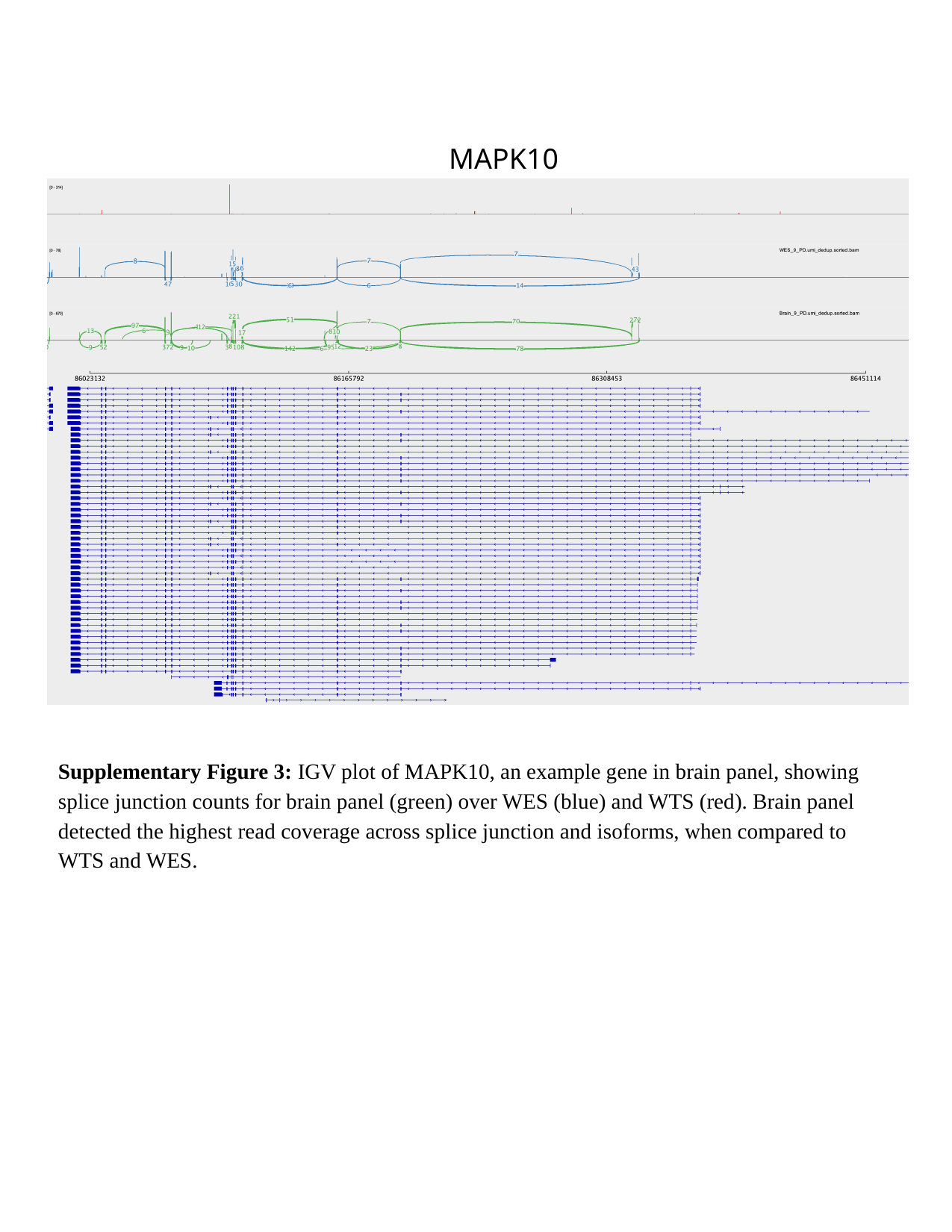

MAPK10
Supplementary Figure 3: IGV plot of MAPK10, an example gene in brain panel, showing splice junction counts for brain panel (green) over WES (blue) and WTS (red). Brain panel detected the highest read coverage across splice junction and isoforms, when compared to WTS and WES.

### Slide 4
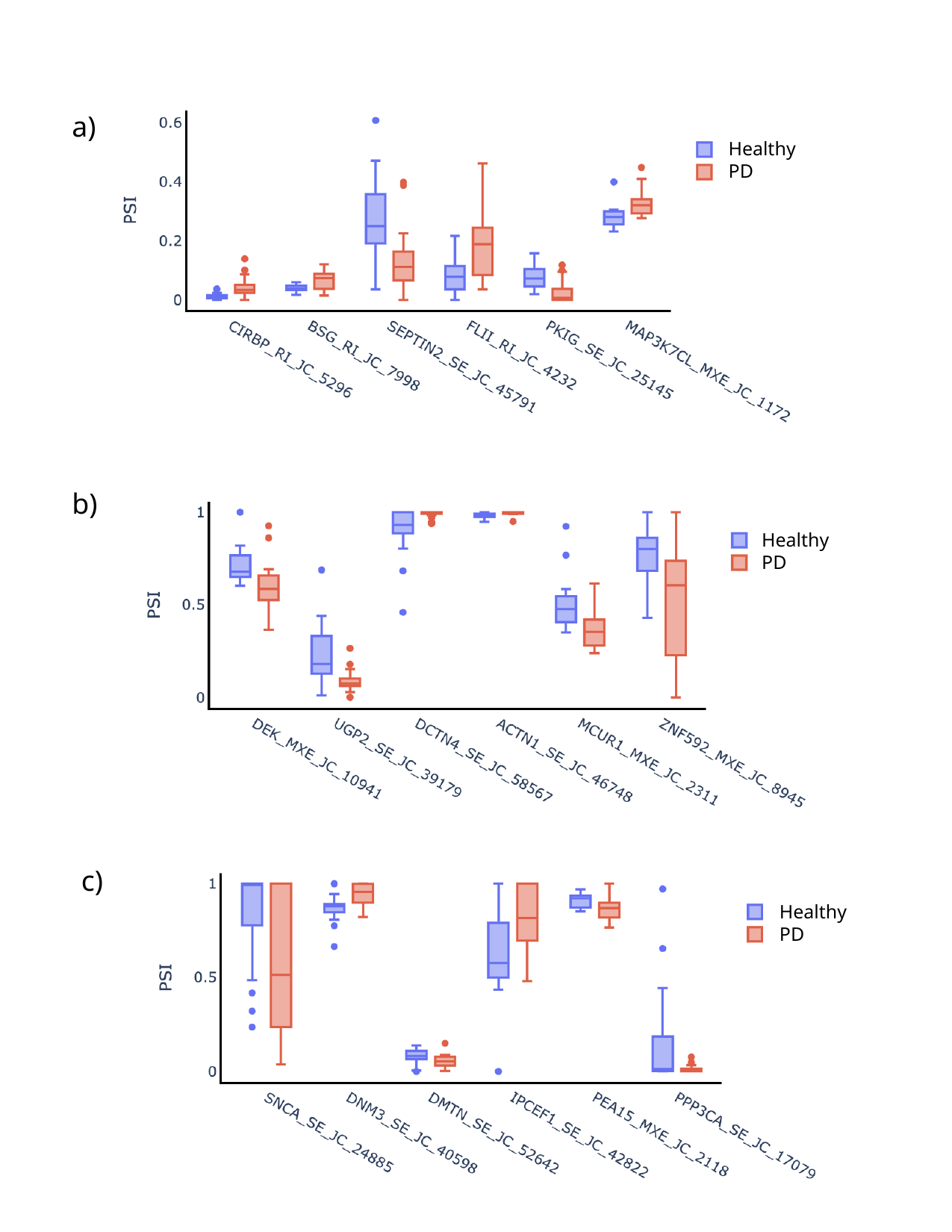

a)
Healthy
PD
b)
Healthy
PD
Healthy
PD
c)
Healthy
PD

### Slide 5
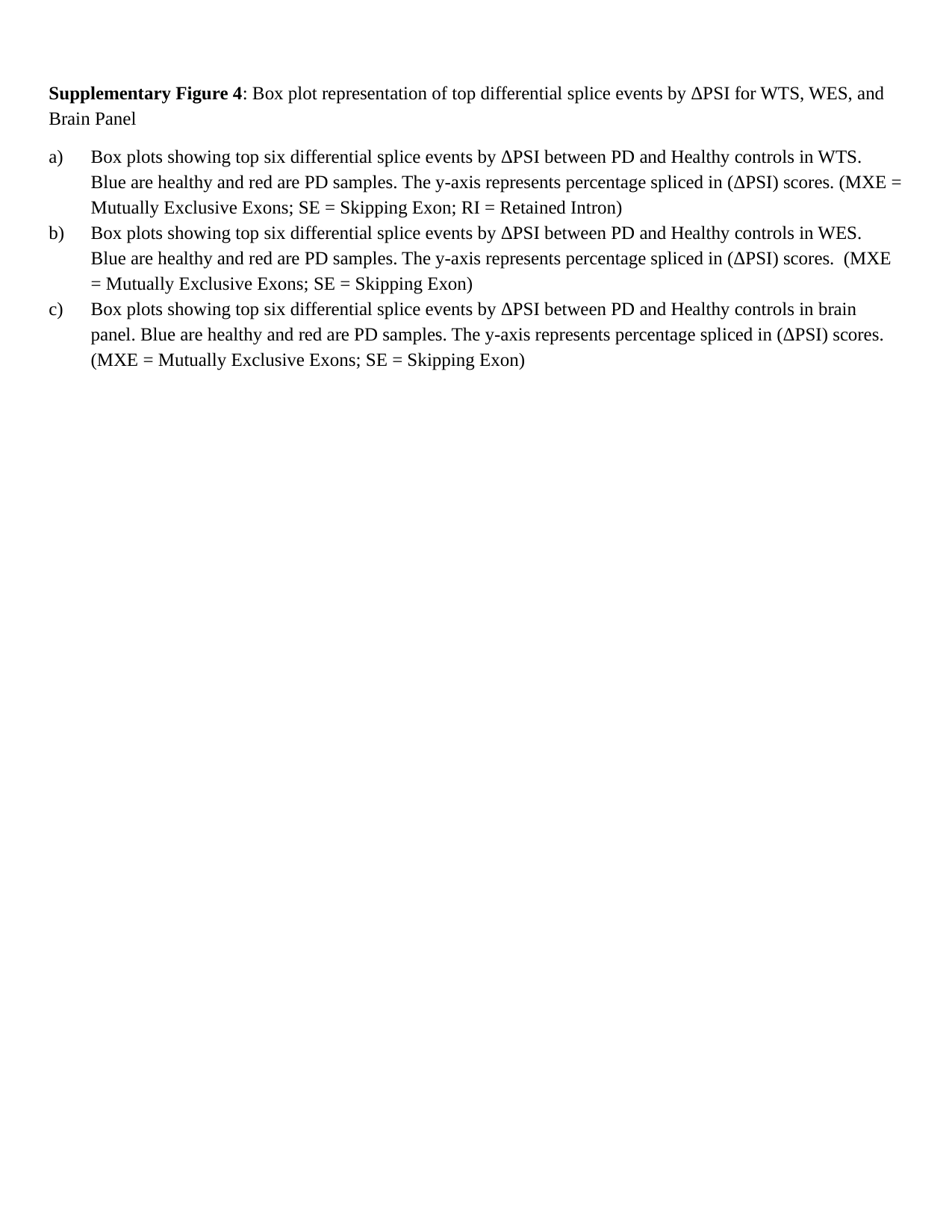

Supplementary Figure 4: Box plot representation of top differential splice events by ΔPSI for WTS, WES, and Brain Panel
Box plots showing top six differential splice events by ΔPSI between PD and Healthy controls in WTS. Blue are healthy and red are PD samples. The y-axis represents percentage spliced in (ΔPSI) scores. (MXE = Mutually Exclusive Exons; SE = Skipping Exon; RI = Retained Intron)
Box plots showing top six differential splice events by ΔPSI between PD and Healthy controls in WES. Blue are healthy and red are PD samples. The y-axis represents percentage spliced in (ΔPSI) scores. (MXE = Mutually Exclusive Exons; SE = Skipping Exon)
Box plots showing top six differential splice events by ΔPSI between PD and Healthy controls in brain panel. Blue are healthy and red are PD samples. The y-axis represents percentage spliced in (ΔPSI) scores. (MXE = Mutually Exclusive Exons; SE = Skipping Exon)

### Slide 6
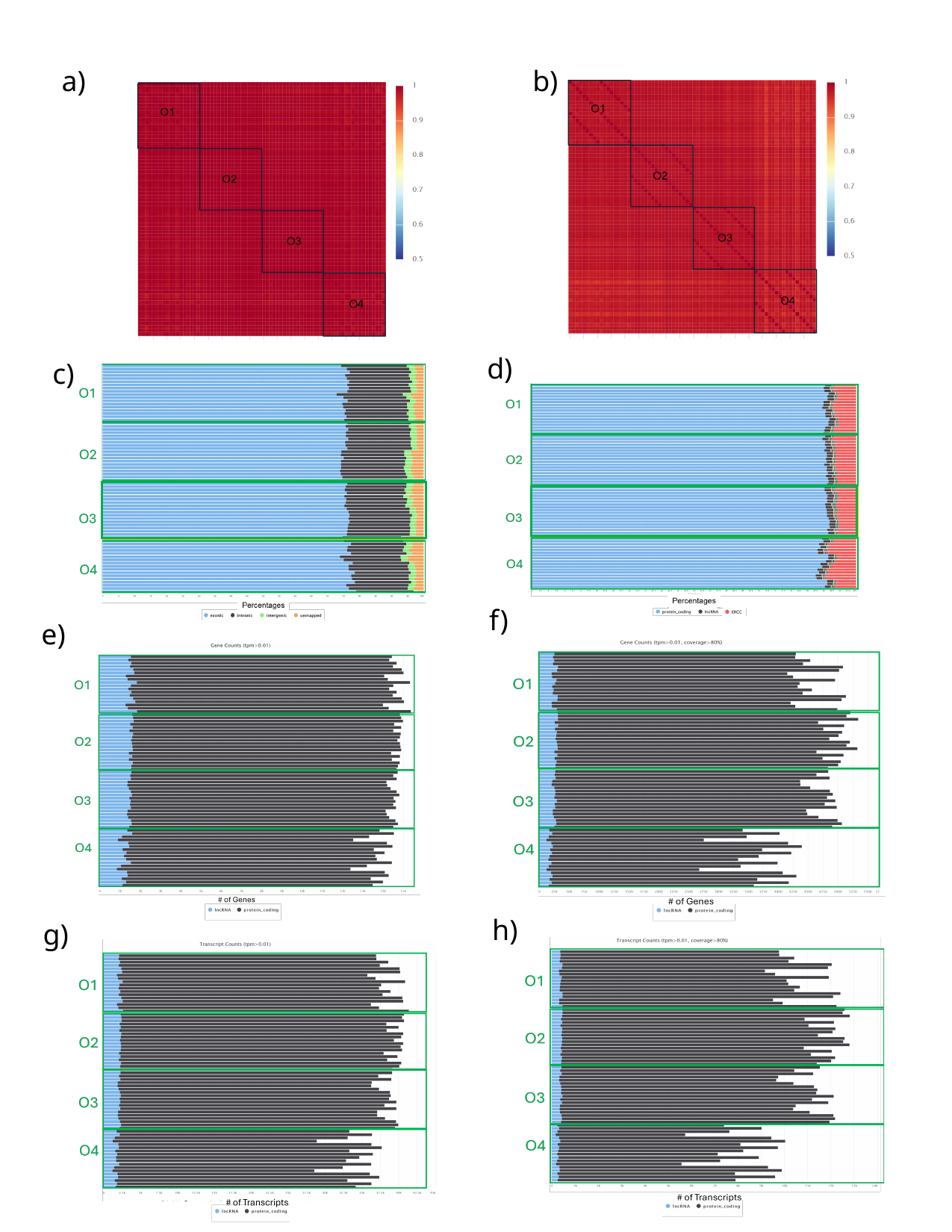

a)
b)
d)
c)
f)
e)
h)
g)

### Slide 7
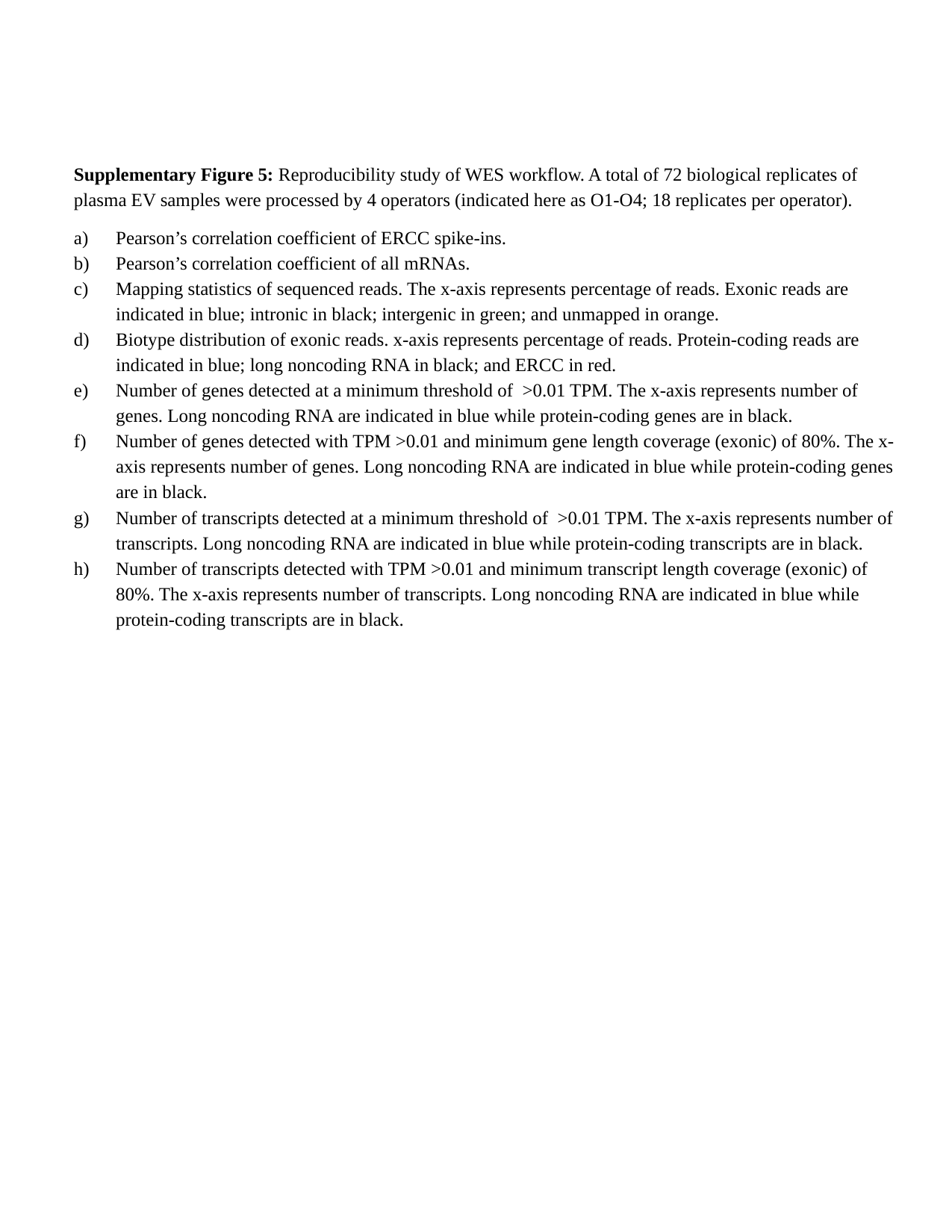

Supplementary Figure 5: Reproducibility study of WES workflow. A total of 72 biological replicates of plasma EV samples were processed by 4 operators (indicated here as O1-O4; 18 replicates per operator).
Pearson’s correlation coefficient of ERCC spike-ins.
Pearson’s correlation coefficient of all mRNAs.
Mapping statistics of sequenced reads. The x-axis represents percentage of reads. Exonic reads are indicated in blue; intronic in black; intergenic in green; and unmapped in orange.
Biotype distribution of exonic reads. x-axis represents percentage of reads. Protein-coding reads are indicated in blue; long noncoding RNA in black; and ERCC in red.
Number of genes detected at a minimum threshold of >0.01 TPM. The x-axis represents number of genes. Long noncoding RNA are indicated in blue while protein-coding genes are in black.
Number of genes detected with TPM >0.01 and minimum gene length coverage (exonic) of 80%. The x-axis represents number of genes. Long noncoding RNA are indicated in blue while protein-coding genes are in black.
Number of transcripts detected at a minimum threshold of >0.01 TPM. The x-axis represents number of transcripts. Long noncoding RNA are indicated in blue while protein-coding transcripts are in black.
Number of transcripts detected with TPM >0.01 and minimum transcript length coverage (exonic) of 80%. The x-axis represents number of transcripts. Long noncoding RNA are indicated in blue while protein-coding transcripts are in black.

### Slide 8
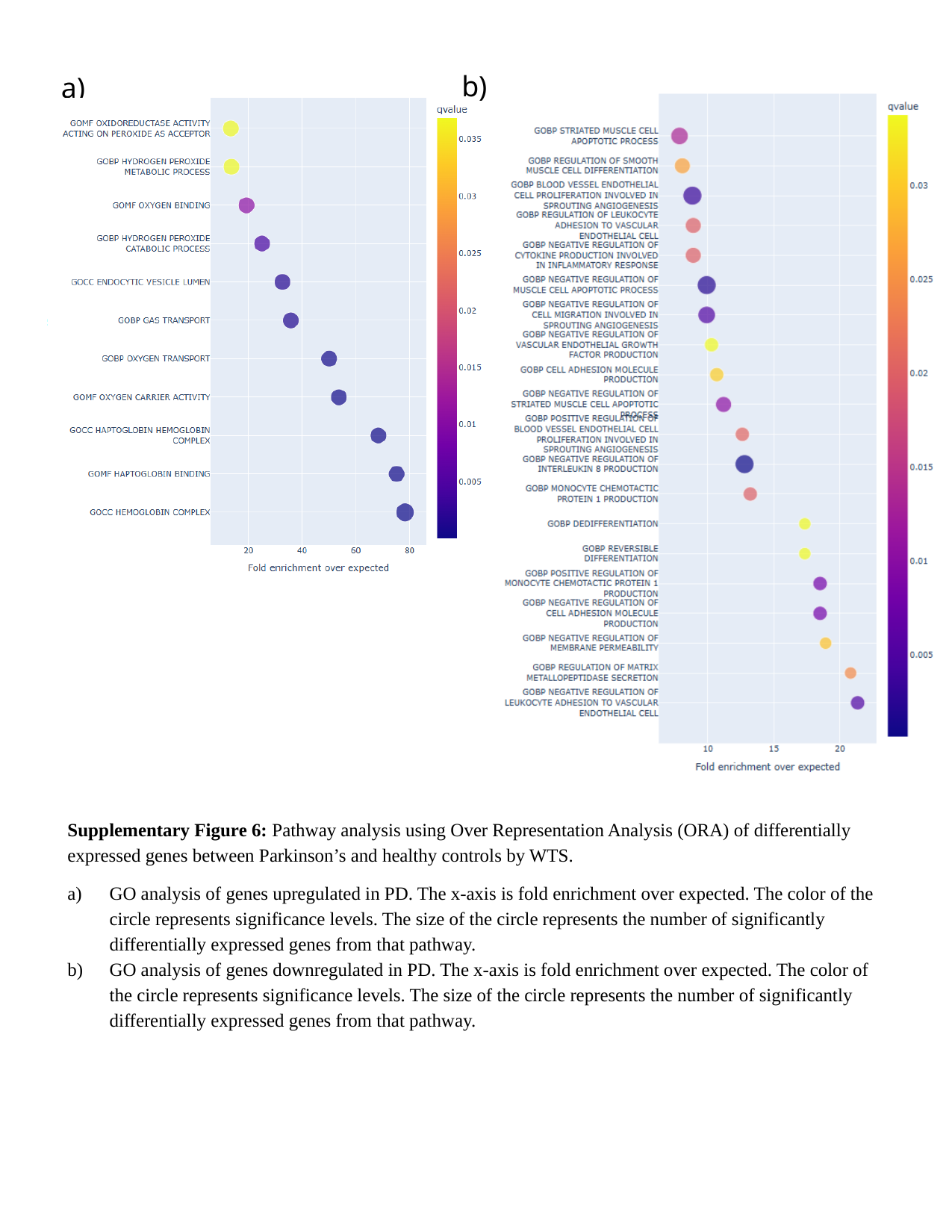

b)
a)
Supplementary Figure 6: Pathway analysis using Over Representation Analysis (ORA) of differentially expressed genes between Parkinson’s and healthy controls by WTS.
GO analysis of genes upregulated in PD. The x-axis is fold enrichment over expected. The color of the circle represents significance levels. The size of the circle represents the number of significantly differentially expressed genes from that pathway.
GO analysis of genes downregulated in PD. The x-axis is fold enrichment over expected. The color of the circle represents significance levels. The size of the circle represents the number of significantly differentially expressed genes from that pathway.

### Slide 9
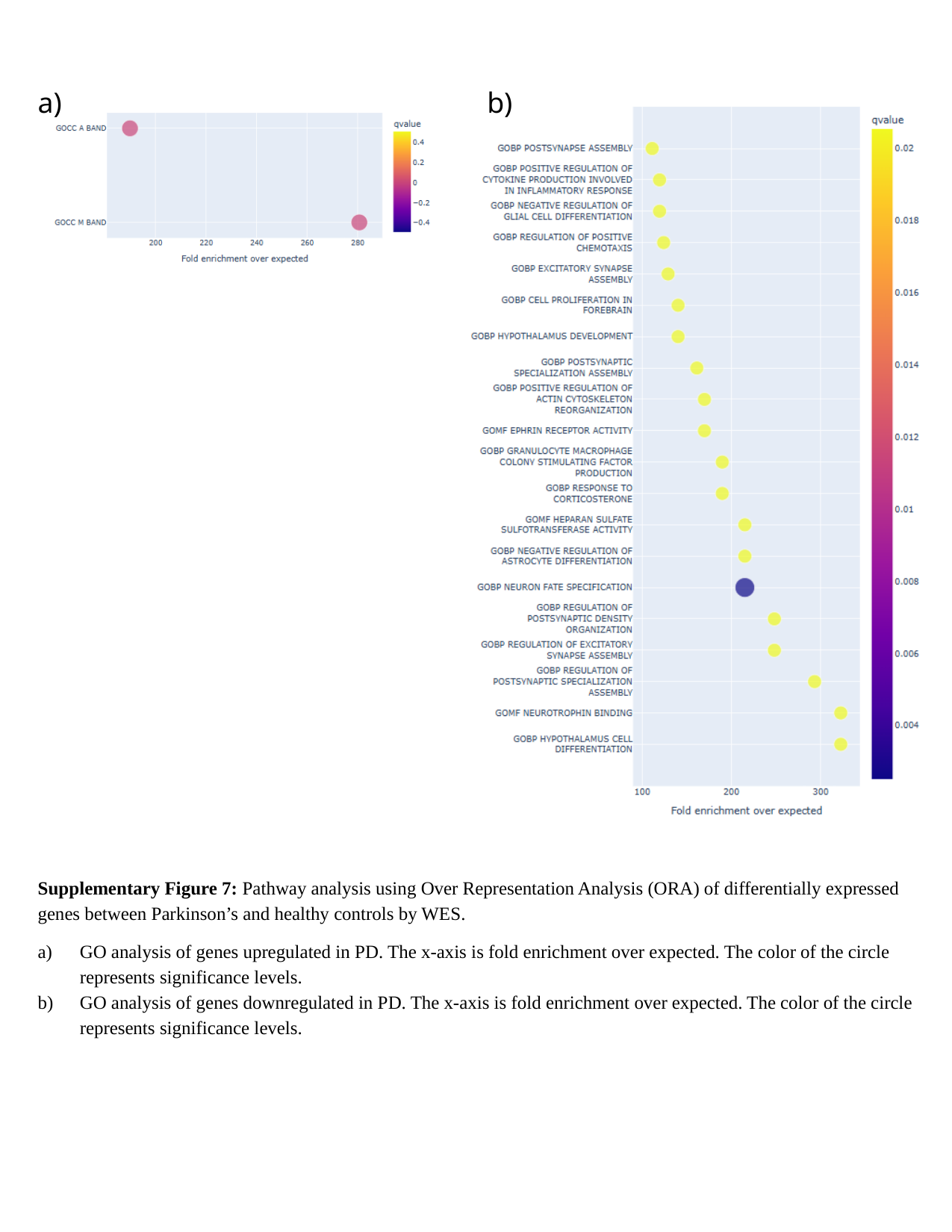

a)
b)
Supplementary Figure 7: Pathway analysis using Over Representation Analysis (ORA) of differentially expressed genes between Parkinson’s and healthy controls by WES.
GO analysis of genes upregulated in PD. The x-axis is fold enrichment over expected. The color of the circle represents significance levels.
GO analysis of genes downregulated in PD. The x-axis is fold enrichment over expected. The color of the circle represents significance levels.

### Slide 10
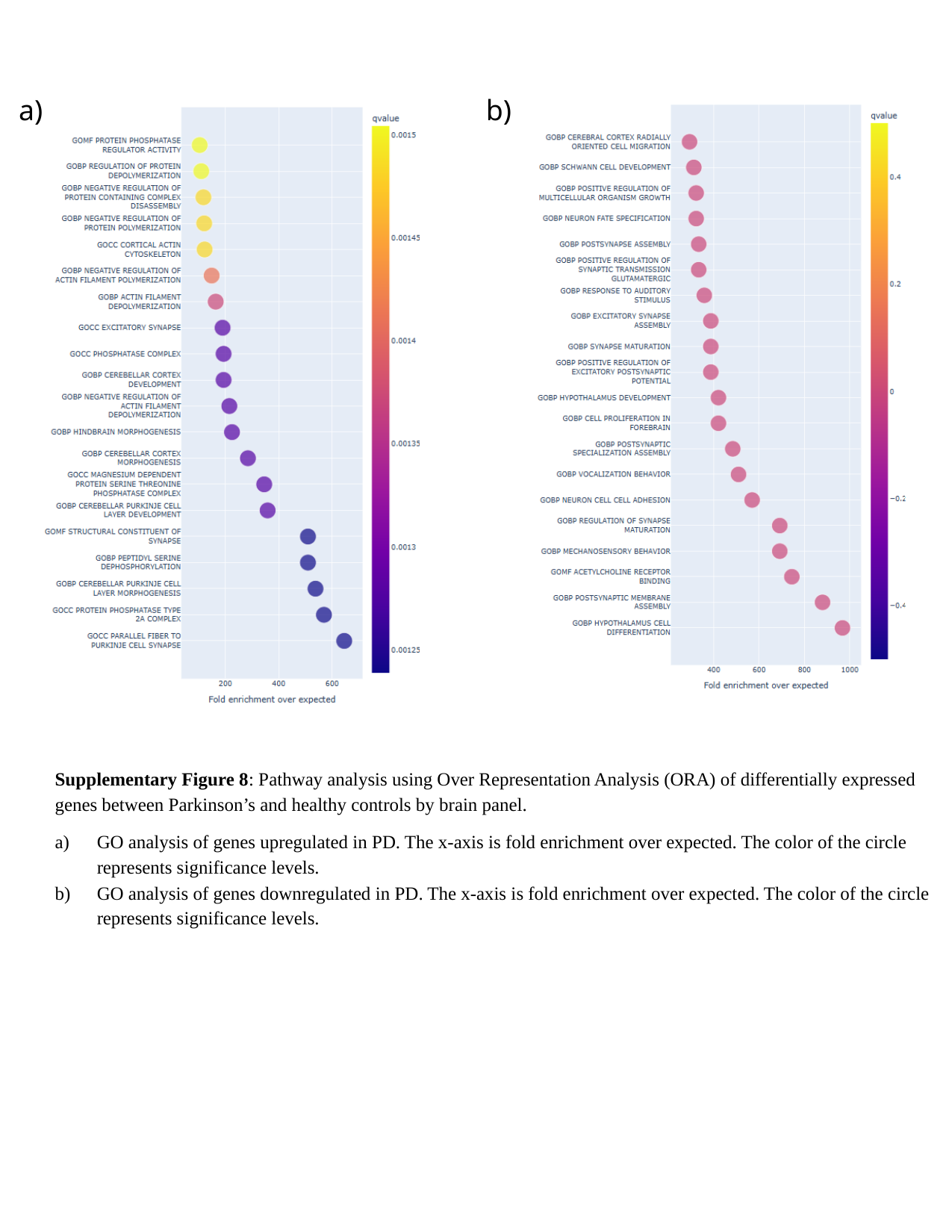

a)
b)
Supplementary Figure 8: Pathway analysis using Over Representation Analysis (ORA) of differentially expressed genes between Parkinson’s and healthy controls by brain panel.
GO analysis of genes upregulated in PD. The x-axis is fold enrichment over expected. The color of the circle represents significance levels.
GO analysis of genes downregulated in PD. The x-axis is fold enrichment over expected. The color of the circle represents significance levels.

### Slide 11
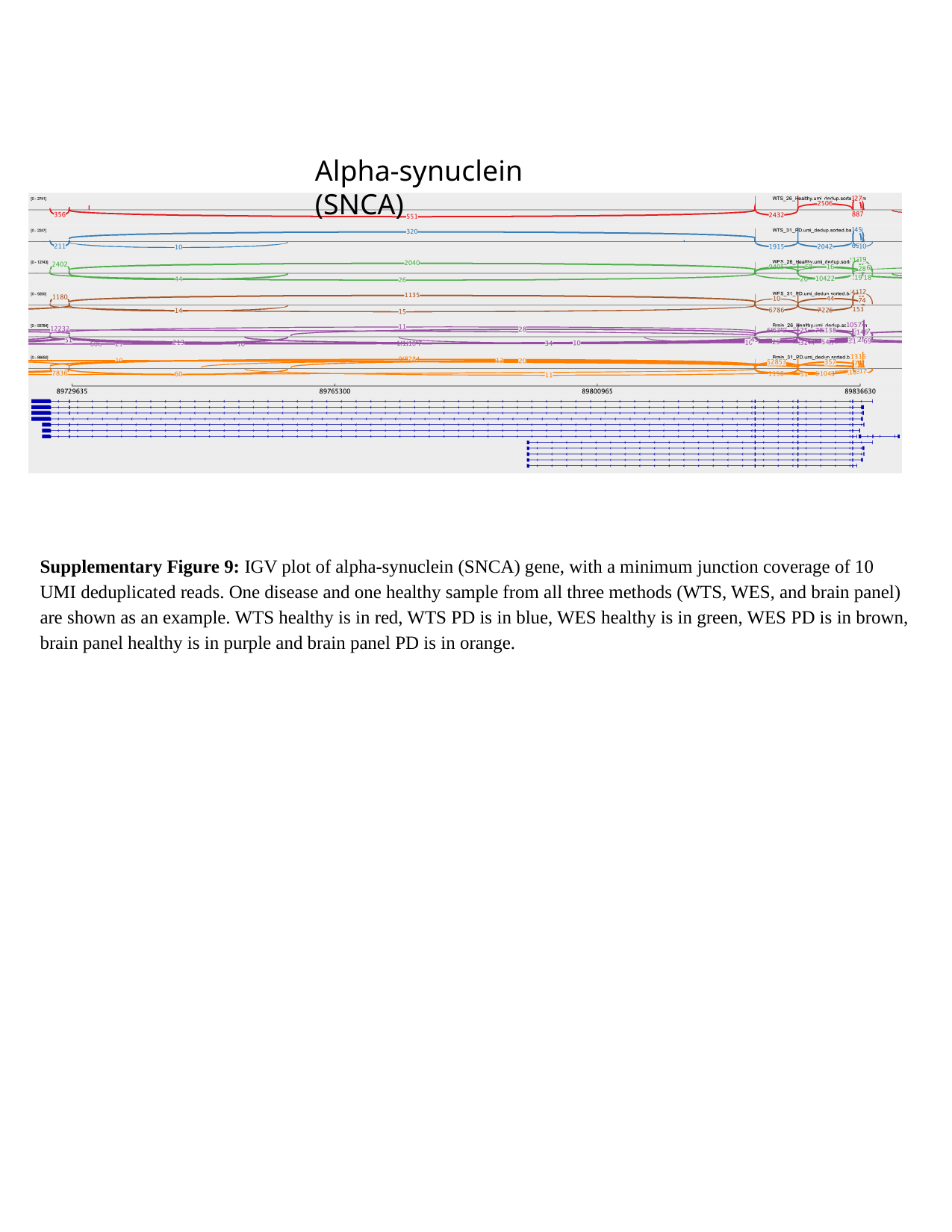

Alpha-synuclein (SNCA)
Supplementary Figure 9: IGV plot of alpha-synuclein (SNCA) gene, with a minimum junction coverage of 10 UMI deduplicated reads. One disease and one healthy sample from all three methods (WTS, WES, and brain panel) are shown as an example. WTS healthy is in red, WTS PD is in blue, WES healthy is in green, WES PD is in brown, brain panel healthy is in purple and brain panel PD is in orange.
