## Supplementary Table Legends for "Deep Profiling of EV Long RNAs Reveals Biofluid-Specific Transcriptomes and Splicing Landscapes"

**Supplementary Tables**

Supplementary Table 1: Table of patient metadata for Parkinson’s disease plasma samples and healthy control plasma samples.

Supplementary Table 2: Table of genes included in the targeted brain panel.

Supplementary Table 3: Table of differentially expressed genes between healthy CSF and healthy plasma in WTS.

Supplementary Table 4: Table of differentially expressed genes between healthy urine and healthy plasma in WTS.

Supplementary Table 5: Table of differentially expressed genes between Parkinson’s Disease and Healthy Plasma in WTS.

Supplementary Table 6: Table of differentially expressed genes between Parkinson’s Disease and Healthy Plasma in WES.

Supplementary Table 7: Table of differentially expressed genes between Parkinson’s Disease and Healthy Plasma in brain panel sequencing.

Supplementary Table 8: Table of differentially expressed splice events between Parkinson’s Disease and Healthy Plasma in WTS.

Supplementary Table 9: Table of differentially expressed splice events between Parkinson’s Disease and Healthy Plasma in WES.

Supplementary Table 10: Table of differentially expressed splice events between Parkinson’s Disease and Healthy Plasma in brain panel sequencing.
